# Mutations in SOD1 induce ALS-related phenotypes in 3D iPSC-derived motor neuron (MN) spheroids

**DOI:** 10.1101/2025.01.28.635126

**Authors:** María José Castellanos-Montiel, Anna Kristyna Franco-Flores, Michael Nicouleau, Ghazal Haghi, Sarah Lépine, Belén Baeza, Carol X.-Q. Chen, Taylor M. Goldsmith, Nathalia Aprahamian, Doris Hua, Mathilde Chaineau, Lale Gursu, Narges Abdian, Eric Deneault, Thomas M. Durcan

## Abstract

A significant challenge in ALS research is the heterogeneity of the disease. Even mutations within the same gene can lead to different disease prognosis. For instance, *in silico* protein modeling predicts distinct properties for distinct SOD1 mutations. With this in mind, in this study, we generated and characterized 3D iPSC-derived MN spheroids carrying homozygous knock-in SOD1 mutations (D90A and G93A), as well as a double mutation (D90A/G93A), to evaluate potential synergistic effects. An isogenic control line with the same genetic background was used for phenotypic comparisons with the knock-in variants. Mutant SOD1 MN spheroids exhibited multiple ALS-related phenotypes including altered SOD1 expression, reduced cell viability, downregulation of neurofilament (NF) subunit expression, hypoactivity, and altered burst activity. Our results highlight the advantages of using 3D MN spheroids as a disease model and stress the importance of considering phenotype variability at the genetic level in ALS.

## Introduction

Amyotrophic lateral sclerosis (ALS) is a heterogeneous, adult-onset neurodegenerative disease characterized by progressive motor neuron (MN) loss in the motor cortex, brainstem, and spinal cord, leading to skeletal muscle weakness and atrophy (1). Following an ALS diagnosis, the average survival is 3 – 4 years, with patients typically succumbing to respiratory failure (2). Sporadic ALS (sALS) accounts for 90% of cases, with the remaining 10%, referred to as familial ALS (fALS), following an inheritance pattern associated with identified gene mutations (3, 4). Mutations in superoxide dismutase 1 (*SOD1*) account for □20% of fALS cases, making it the second most common genetic cause of the disease, surpassed only by C9orf72 mutations, which account for □40% of fALS cases (4, 5).

SOD1 is a 32 kDa homodimeric enzyme whose primary function is to mitigate oxidative stress by converting free superoxide radicals into oxygen and hydrogen peroxide (6, 7). Early evidence from post-mortem brain tissue of sALS and fALS patients suggests that loss-of-function mechanisms resulting from SOD1 haploinsufficiency contribute to increased oxidative stress levels in neurons (8). However, later evidence has shown a limited correlation between enzyme activity and the onset or clinical forms of the disease (9). In turn, it was found that misfolded mutant SOD1 can aggregate and acquire toxic gain of function properties (8, 10). Nevertheless, *in silico* protein modeling and *in vitro* aggregation propensity studies have shown that these aggregation properties can differ between SOD1 variants (11, 12). Thus, a central question in the ALS field is whether SOD1 mutations result in a loss of function, a gain of toxic function, or both.

In 1994, Gurney and colleagues introduced the first transgenic mouse model for ALS, which overexpresses the human SOD1 gene with a glycine-to-alanine mutation at position 93 (SOD1^G93A^) (13, 14). To date, the SOD1^G93A^ mouse model remains the gold standard for motor deficit assessment in preclinical studies (15). However, issues arising from overexpression, inconsistencies in disease progression, and other limitations cast doubt on the complete translatability of this model to the human context (15). Induced pluripotent stem cells (iPSCs) provide access to human neurons derived from patients with neurodegenerative diseases, including MNs, enabling the study of neurodegeneration *in vitro*. Currently, patient iPSC-derived MNs are used alongside animal models to study ALS (16, 17). Moreover, the advent of CRISPR/Cas9 genome editing technology enables the creation of isogenic controls for accurate phenotypic comparisons (18, 19).

Various protocols exist for differentiating iPSC-derived MNs, typically resulting in a 2D monolayer of neurons (20). However, this approach poses technical challenges, as MNs tend to aggregate and detach over time, limiting its suitability for long-term analysis. This is particularly important when modeling neurodegenerative diseases, where disease phenotypes may only emerge after extended culture periods. (21, 22). In a study from our group, we successfully differentiated 2D iPSC-derived MNs harboring ALS-related mutations and maintained them in culture for □42 days under optimal conditions for analysis (18). Nevertheless, the lack of a significant reduction in MN viability in the ALS lines implies that the MNs may not have been mature enough to detect a disease phenotype at the examined time points, highlighting the challenges associated with long-term culture and emphasizing the need for strategies to overcome these limitations. Advancements in 3D *in vitro* systems, such as organoids and spheroids, provide environments that closely mimic cellular development, bringing us closer to replicating human physiology and pathology (23, 24, 25, 26). Furthermore, 3D cultures overcome the detachment issues seen in 2D cultures, enabling extended analysis.

In this study, we observed a number of ALS-related phenotypes in 3D iPSC-derived MN spheroids (23) generated from iPSCs carrying homozygous knock-in mutations at positions D90A and G93A, as well as in a double mutant SOD1^D90A/G93A-KI^ iPSC line developed previously (19). We have previously demonstrated that MNs can be cultured for up to 28 days while maintaining the expression of well-established MN markers (23). Here, we extended the culture period to 56 days to determine whether the observed phenotypes become more pronounced over longer durations. To ensure the accuracy of our findings, we compared the phenotypic changes in these mutant lines to MN spheroids derived from the isogenic control iPSC line (27), which shares the same genetic background except for the introduced *SOD1* mutations. This approach provides a 3D scalable model for investigating ALS distinct cellular pathologies associated with SOD1 mutations.

## Materials and methods

### Generation and maintenance of human iPSCs

Following a streamlined CRISPR/Cas9 workflow, the validated non-carrier AIW002-02 iPSC line (27) was used to engineer a homozygous double knock-in iPSC line carrying two *SOD1* mutations known to encode fALS variants: D90A and G93A (SOD1^D90A/G93A-KI^) (19). Here, we describe the correction of the SOD1^D90A/G93A-KI^ iPSC line to generate two new iPSC lines, each carrying a single homozygous mutation, either D90A (SOD1^D90A-KI^) or G93A (SOD1^G93A-KI^) (**Additional file 1**) (**Additional files 2-4: Tables S1-3**) (**Additional file 8**). For this study, the SOD1^D90A-KI^, SOD1^G93A-KI^, and SOD1^D90A/G93A-KI^ iPSC lines were used alongside the original AIW002-02 iPSC line as the isogenic control for precise phenotypic comparisons. Briefly, all iPSC lines were maintained on Matrigel (#354277, Corning)-coated dishes in mTeSR1^TM^ medium (#85850, STEMCELL Technologies) and passaged at 80% confluence using Gentle Cell Dissociation Reagent (#100-0485, STEMCELL Technologies) (27). The complete profiles of the iPSCs, detailed culture procedures, and methods for quality control analyses– including immunocytochemistry for pluripotency associated markers, qPCR to assess chromosome copy numbers, karyotyping and virology testing–are available in (19, 27) (**Additional file 1**) (**Additional file 9**). Prior to starting the experiments, iPSCs were tested weekly and found to be free from mycoplasma. The use of iPSCs in this research was approved by the McGill Research Ethics Board (IRB Study Number A03-M19-22A).

### Generation and characterization of iPSC-derived motor neuron (MN) spheroids

For each iPSC line (AIW002-02, SOD1^D90A-KI^, SOD1^G93A-KI^, and SOD1^D90A/G93A-KI^), three independent differentiations were performed to obtain motor neuron progenitor cells as previously described (MNPCs) (**Supplementary Figure 3**) (18, 19, 23, 28). Next, MNPCs were used to generate and characterize MN spheroids following a published optimized workflow (23). Briefly, MNPCs were dissociated into a single cell suspension using Accutase (#07922, Thermo Fisher Scientific) and plated into round-bottom ultra-low-attachment 96-well plates (#707, Corning) to promote their aggregation as 5K or 10K MN spheroids that were assessed at different time points at the morphological, transcript, protein, and functional level.

### Size profiling

Bright-field images of 5K or 10K MN spheroids were acquired with an EVOS^TM^ XL Core (Thermo Fisher Scientific) light microscope after 14, 28, and 56 days in MN induction and maturation medium. System 20X/0.40 objective. Image size, 1024 x 768 pixels. Pixel size, 0.449 µm x 0.449 µm. The longitudinal growth of each MN spheroid was followed by saving their location within the plate. Next, a CellProfiler pipeline available online was used to measure the area of each MN spheroid in pixels (https://doi.org/10.17605/OSF.IO/V84WS) (accessed on 9 September 2024) (23). Finally, the area in pixels was manually transformed into μm^2^ by multiplying x 0.449 µm x 0.449 µm. For each cell line, twenty MN spheroids from each of three different batches were analyzed.

### Clearing and immunocytochemistry

5K MN spheroids were fixed, cleared, and immunostained at 14, 28 and 56 days, as previously described with minor modifications (23). During the Clear Unobstructed Brain/Body Imaging Cocktail and Computational Analysis (CUBIC) protocol (23, 29), 56-day-old MN spheroids required a 72 h treatment with CUBIC R1 due to their size. Images of the immunostained MN spheroids were acquired with the Opera Phenix High-Content Screening System using the PreScan function to find the spheres within the focal plane at 5X and then perform the imaging at 20X. System 5X/0.16 and 20Xwater/1.0 objectives. Image size 512 × 512 pixels. Voxel size, 0.29 µm × 0.29 µm × 5 µm. Data was extracted to be organized and analyzed by an in-house script developed in MATLAB. Images were analyzed as raw Z-stacks without altering brightness and contrast. A list of all primary and secondary antibodies used is provided in **Additional file 5 and 6: Tables S4 and S5**.

### RT-qPCR

For each condition, sixty 5K MN spheroids were pooled into a 1.7 mL collection tube (#87003-294, VWR) and total RNA was isolated using the miRNeasy Micro Kit (#217084, Qiagen) according to the manufacturer’s instructions. Subsequently, reverse transcription reactions for cDNA synthesis, followed by qPCR reactions, were performed as previously described (23). The mean between Actβ and GAPDH was used as the endogenous control for normalization. A list of all the TaqMan probes used is provided in **Additional file 7: Table S6.**

### Western Blotting

Sixty 10K MN spheroids were pooled into a single 1.7 mL collection tube for whole protein extraction. Briefly, samples were incubated with cold 1X RIPA buffer (#20-188, Millipore) containing a cocktail of protease (#11697498001, Roche) and phosphatase inhibitors (#04906837001, Roche) for 15 minutes at 4°C. Every 5 minutes, the samples were mechanically dissociated using a pipette. Finally, the samples were centrifuged at 10,000 g for 30 min at 4°C, and supernatants were collected for protein quantification.

Protein concentration in the supernatants was quantified using the DC Protein Assay (#5000116, Bio-Rad) according to the manufacturer’s instructions. For blotting NFL, NFM, and NFH, 1 µg of protein was loaded onto a 7.5% SDS-PAGE gel and run at 70□V for 15 min, then at 120□V for ∼1.5 h. Semi-dry transfer to nitrocellulose membranes was performed using the Trans-Blot Turbo Transfer System (Bio-Rad) for 10 min at 2.5 A, up to 25□V. For blotting βIII-tubulin, SOD1, and cleaved caspase-3, 1 µg (or 5 µg for cleaved caspase-3) of protein was loaded onto a 12% SDS-PAGE gel, run at 70□V for 15 min, then at 120□V for ∼1 h. Semi-dry transfer to nitrocellulose membranes was performed for 30 min up to 1.0 A, at 25□V. After transfer, the membranes were blocked with 5% BSA (or 5% skimmed milk [#SKI400.1, Bioshop] for cleaved caspase-3) in 1X TBS buffer containing 0.1% Tween 20 (#TWN510, Bioshop) (blocking solution) for 1□h at room temperature (RT) with continuous shaking, followed by overnight incubation with primary antibodies, diluted in their respective blocking solutions, at 4°C with continuous shaking. After three 10-min washes with 1X TBS containing 0.1% Tween 20 (washing solution) and continuous shaking, membranes were incubated with horseradish peroxidase (HRP)-conjugated antibodies, diluted in their respective blocking solutions, for 2□h at RT with continuous shaking. After three 10-min washes with washing solution and continuous shaking, protein bands were detected using the Clarity^TM^ Western ECL (#1705060, Bio-Rad) (or Clarity^TM^ Max Western ECL [#1705062S, Bio-Rad] for cleaved caspase-3) according to the manufacturer’s instructions and visualized using a Chemidoc MP Imaging System (Bio-Rad). Quantification was performed with the Image Lab 6.0.1 software (Bio-Rad), using β-Actin and GAPDH as loading controls. A list of all antibodies used is provided in **Additional file 5 and 6: Tables S4 and S5**.

### CellTiter-Glo® 3D Cell Viability Assay

10K MN spheroids were grown for 56 days in round-bottom ultra-low-attachment 96- well plates. At the endpoint and before starting the experiment, bright-field images of the MN spheroids were acquired with an EVOS^TM^ XL Core light microscope for normalization of the luminescence measures against MN size. System 20X/0.40 objective. Image size, 1024 x 768 pixels. Pixel size, 0.449 µm x 0.449 µm. Each MN spheroid along with 100 µL of medium was transferred into a well of a white round-bottom 96-well plate (#3789A, Corning) using wide-orifice low-binding tips and the CellTiter-Glo® 3D Cell Viability Assay (#G9682, Promega) was performed following the manufacturer’s instructions. Luminescence readings as random luminescence units (RLU) were acquired using a SpectraMax iD3 multi-mode microplate reader (Molecular Devices). After RLU acquisition, each measurement was normalized against the area in pixels of its corresponding MN spheroid to obtain a ratio (RLU/area in pixels). For comparisons across cell lines, ratios were converted to percentages, with the ratio of the isogenic control set as 100% survival.

### Detection of NFL release in cell culture medium

10 K MN spheroids were cultured for 56 days in round-bottom ultra-low-attachment 96- well plates, with no medium changes throughout the entire culture period. Media from five MN spheroids (60 µL per spheroid) were pooled into a 1.7 mL collection tube for each condition, resulting in a final volume of 300 µL, and were snap-frozen in liquid nitrogen. For the experiment, all samples were thawed and centrifuged at 1000 g for 20 min at 4°C. The assay was then performed according to the manufacturer’s instructions (#NBP2-81184, Novus Biologicals).

### Microelectrode array (MEA) recordings

5K MN spheroids were generated into round-bottom ultra-low-attachment 96-well plates. After 7 days in culture, a maximum of six MN spheroids were pooled into a 0.6 mL collection tube and plated into an individual well of a Cytoview 24–well MEA plate (#M384-Tmea-24w, Axion Biosystems) using wide-orifice low-binding tips as previously described (23). Cytoview 24-well MEA plates were coated with a 10 μg/mL solution of poly-L-ornithine (PLO, #P3655, Sigma-Aldrich) diluted in 1X PBS. After 24 h at 37 °C, PLO was washed out by performing three washes with 1X PBS. Immediately afterward, MEA plates were coated with a 5 μg/mL solution of laminin (#L2020, Sigma-Aldrich) diluted in DMEM/F-12 (#10565018, Thermo Fisher Scientific) and incubated for 24 h at 37 °C.

MEA recordings were performed on days 14, 28, 42, and 56 post-plating (p-p) the MN spheroids. Before every recording, MN induction and maturation medium was replaced with 1X artificial cerebrospinal fluid (aCSF) prepared as previously described (23). Osmolarity (□305mOsm/L) and pH (□7.4) were monitored before adding the aCSF to the MN spheroids to ensure physiological values. MEA plates with 1X aCSF were kept in the incubator for 1 h before being transferred to the Axion Maestro Edge (Axion Biosystems) and allowed to equilibrate at 37°C and 5% CO2 for 5 min before recording. Data was collected for 5 min using the Axis Navigator software (provided by Axion Biosystems, version 1.5.1.12, Atlanta, GA, USA). A band-pass filter of 3 kHz (low-pass) to 200□Hz (high-pass) was applied. After recording, the MEA plate was removed from the instrument and the 1X aCSF was replaced with MN induction and maturation medium to keep the cells in culture for the following recording points. At day 56 p-p, after the initial 5-min recording in 1X aCSF, the plates were removed from the instrument and dosed with either vehicle (H_2_O) or 1 mM tetrodotoxin (TTX, # T8024, Sigma-Aldrich) before performing a second 5-min recording. This was the endpoint of the MEA recordings. For analysis, a “spike” was determined as a short extracellular electrical event with a peak voltage six times or greater than the standard deviation of the estimated “noise” signal. A “burst” was determined as ≥ 5 spikes with no more than 100 ms separating each spike. Only data from MN spheroids that covered 90–100% of the electrodes and had four consecutive recording time points were included in the analysis.

## Results

### iPSCs carrying the SOD1^D90A-KI^, SOD1^G93A-KI^, and SOD1^D90A/G93A-KI^ mutations were successfully differentiated into 2D motor neuron progenitor cells (MNPCs)

SOD1^D90A-KI^, SOD1^G93A-KI^, SOD1^D90A/G93A-KI^, and AIW002-02 (isogenic control) iPSCs expressing the pluripotency-associated markers homeobox protein NANOG, podocalyxin-like protein 1 (Tra-1-60), stage-specific embryonic antigen 4 (SSEA-4), and octamer-binding transcription factor 4 (OCT4) (**Additional file 9: Supplementary Figure 2A/B**) were differentiated into iPSC-derived motor neuron progenitor cells (MNPCs). The cellular identity of the MNPCs from all the lines was characterized by immunocytochemistry and qPCR, at the same passage number. To determine the dorsoventral identity of the MNPCs, we quantified the expression levels of PAX6 and OLIG2, as these markers are expected to be elevated in the MN progenitor domain (pMN) of the spinal cord (30). *OLIG2* was found at comparable levels between cell lines at the transcriptional level, while *PAX6* was downregulated in SOD1^D90A/G93A-KI^ MNPCs (p = 0.023) but reached similar levels to the isogenic control at the MN stage (**Additional file 10: Supplementary Figure 3A**). In addition, the percentage of cells positive for the double immunostaining PAX6^+^/OLIG2^+^ confirmed a comparable expression between cell lines at the protein level (AIW002-02, 43.1 ± 4.09; SOD1^D90A-KI^, 55.97 ± 6.87; SOD1^G93A-KI^, 49.33 ± 4.84; SOD1^D90A/G93A-KI^, 47.3 ± 14.38) (**Additional file 10: Supplementary Figure 3B-G**). Finally, the transcriptional expression of the marker *NKX6.1* further confirmed the ventralization of the MNPCs from all cell lines (**Additional file 11: Supplementary Figure 4A-D**) (30).

The rostrocaudal identity of MNs is defined by the differential expression of several HOX genes in their progenitor cells along the neural tube during early embryonic development (31). Thus, we analyzed the expressions of *HOXA5*, *HOXB8*, and *FOXG1* as markers of brachial, thoracic, and forebrain MNs, respectively. MNPCs from all the cell lines show almost no transcriptional expression of *FOXG1*, while expressing higher levels of *HOXB8* compared to *HOXA5*, suggesting that they possess a brachial-thoracic identity (**Additional file 11: Supplementary Figure 4A-D**).

MNPCs generated from three independent iPSC inductions per cell line exhibited comparable expression of key MNPC markers (PAX6 and OLIG2) at both the transcript and protein levels. Additionally, the enriched expression of HOXB8, relative to FOXG1, confirmed their spinal identity. Thus, after characterization, we concluded that all MNPCs were comparable, and we proceeded with the generation of MN spheroids from the four cell lines that were assessed at multiple levels at 14, 28, and 56 days in culture.

### iPSC-derived MN spheroids from SOD1 mutants display size differences compared to the isogenic control

The morphology assessment of the MN spheroids generated from the four different lines (**Figure 1A**) showed that SOD^G93A-KI^ and SOD1^D90A/G93A-KI^ MN spheroids were greater in size than AIW002-02 MN spheroids at 14, 28, and 56 days (**Figure 1B-E**). In contrast, SOD1^D90A-KI^ MN spheroids were smaller in size than AIW002-02 MN spheroids across all time points (**Figure 1B-E**). To assess if the disparity in size was linked to differences in the number of proliferative cells within the MN spheroids, we performed qPCR and immunocytochemistry analyses to detect the expression of *SOX1*.

**Figure 1.**
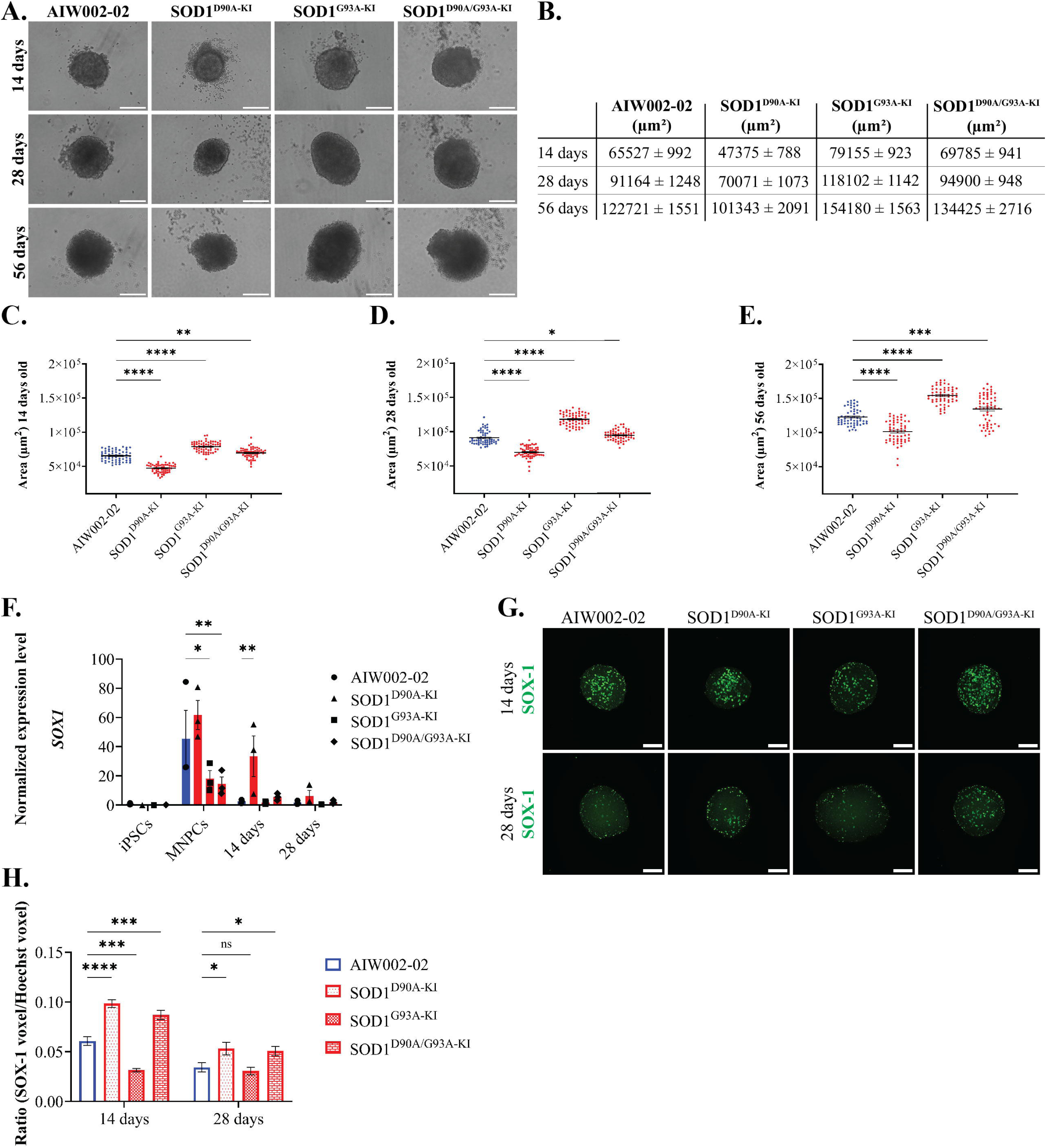
Size profiling using a CellProfiler pipeline reveals size variations in SOD1 MN spheroids. **A.** Bright-field images of AIW002-02 (isogenic control), SOD1^D90A-KI^, SOD1^G93A-KI^, and SOD1^D90A-KI/G93A-KI^ MN spheroids were acquired after 14, 28 and 56 days in culture. Scale bar, 200 µm. **B.** Summary of the mean MN spheroid size for each cell line at 14, 28, and 56 days. The size of the MN spheroids was determined from the bright-field images using a CellProfiler pipeline. **C-E.** Scatter plots showing the distribution of ninety MN spheroids analyzed per cell line at 14, 28, and 56 days. Data shown as mean ± SEM, N=3, n=30. **F. A** RT-qPCR targeting *SOX1* was performed to assess whether the size differences of the MN spheroids were linked to changes in the number of proliferative cells. Data shown as mean ± SEM, N=3, n=3. **G-H.** In addition, SOX-1 (green) expression was analyzed by immunocytochemistry to examine its correlation to *SOX1* mRNA levels. Images represent the maximal projection of 50 optical slices acquired through confocal imaging. Scale bar, 100 µm. Data shown as mean ± SEM, N=3, n=4. Significance was determined using a two-way ANOVA, followed by a post hoc Dunnett’s test using AIW002-02 as the reference sample. *, p ≤ 0.05; **, p ≤ 0.01; ***, p ≤ 0.001; ****, p ≤ 0.0001; ns, non-significant (p > 0.05).

*SOX1* (**Figure 1F**), encoding SOX-1 (**Figure 1G/H**), a marker associated with pan-neuronal proliferating precursors (32), was mostly absent in all cell lines at the iPSC stage. At the MNPC stage, *SOX1* expression was downregulated in the SOD1^G93A-KI^ and SOD1^D90A/G93A-KI^ cell lines. However, *SOX1* expression reached comparable levels to the isogenic control in SOD1^G93A-KI^ and SOD1^D90A/G93A-KI^ MN spheroids after 14 and 28 days. In line with this, SOD1^G93A-KI^ MN spheroids matched the isogenic control at the protein level (SOX-1) at 28 days. Interestingly, while SOD1^D90A/G93A-KI^ MN spheroids do not exhibit significant changes in *SOX1* after 14 and 28 days, there is an upregulation of SOX-1 after 14 days that remains until 28 days. Similarly, SOD1^D90A-KI^ MN spheroids showed an increased expression of *SOX1* that was also significant at the protein level (SOX-1) after 14 and 28 days.

These results suggest that the larger size of SOD1^D90A/G93A-KI^ MN spheroids could be attributed to a slower decline in the population of proliferative neural precursors expressing SOX-1. However, SOD1^D90A-KI^ MN spheroids showed the highest levels of SOX-1 but were smaller than the isogenic control, while SOD1^G93A-KI^ MN spheroids were larger than the isogenic control despite having similar SOX-1 decline patterns. Taken together, these results indicate that the size difference observed for the different MN spheroids is not driven by the rate of neural precursor population decline.

### SOD1 MN spheroids retain a cellular identity comparable to their isogenic control

To determine if the proportion of differentiated cells was similar within MN spheroids regardless of their size, we assessed the transcriptional and protein profiles of AIW002-02, SOD1^D90A-KI^, SOD1^G93A-KI^, and SOD1^D90A-KI/G93A-KI^ MN spheroids by qPCR and immunocytochemistry respectively.

The transient co-expression of insulin gene enhancer 1 (ISL1) and MN and pancreas homeobox 1 (MNX1, better known as HB9) can be used to identify nascent lower motor neurons (LMNs) from the brainstem and spinal cord (20, 33). However, medial and lateral LMNs belonging to the lateral motor column, which innervates limb muscles, are expected to downregulate both ISL1 and HB9 as they mature (34, 35). Thus, the expression of choline acetyltransferase (ChAT) is also analyzed to identify more mature LMNs (20, 34). With this in mind, the expressions of ISL1, HB9 and CHAT were assessed in 28 and 56-day-old MN spheroids at both the transcript and protein levels (**Additional file 12: Supplementary Figure 5A**) (**Figure 2**).

**Figure 2.**
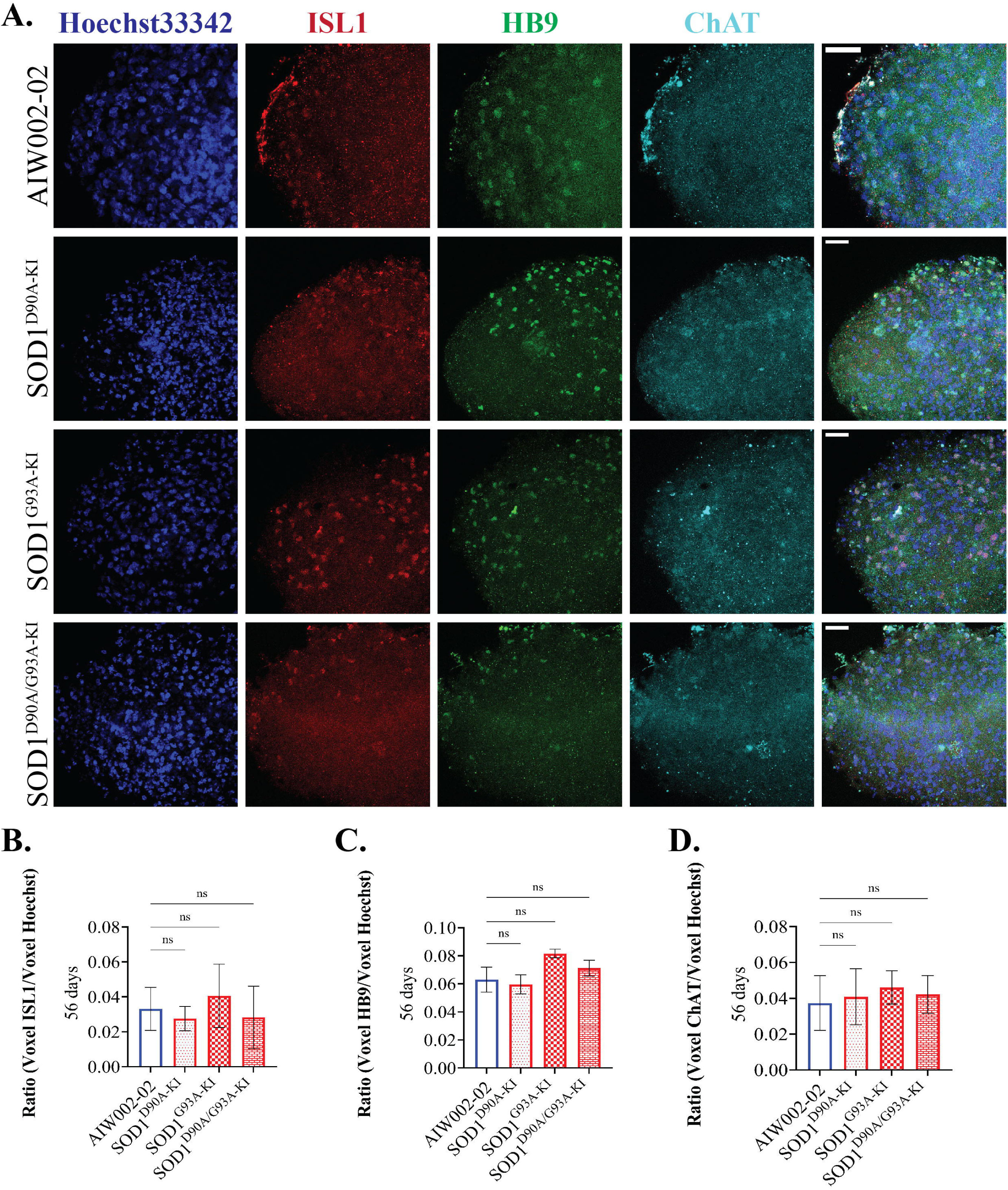
MN markers are consistently expressed at the transcript and protein levels in MN spheroids from the different lines. **A.** Immunocytochemistry analyses on 56-day-old MN spheroids were conducted to assess whether the protein levels of ISL1 (red), HB9 (green) and, ChAT (cyan) correlated to their mRNA expression levels. Images show a zoom-in of the maximal intensity projection derived from 12 optical slices acquired through confocal imaging. Scale bar, 50 µm. **B-D.** Quantification revealed no significant differences in the protein levels of ISL1, HB9, and ChAT across the different cell lines. Data shown as mean ± SEM, N=3, n=4. Significance was determined using a two-way ANOVA, followed by a post hoc Dunnett’s test using AIW002-02 as the reference sample. *, p ≤ 0.05; ****, p ≤ 0.0001; ns, non-significant (p > 0.05).

At the transcriptional level, *ISL1* and *HB9* are completely absent at the iPSC stage. As the cells are induced to MNPCs and later differentiated into MNs, both genes upregulate their expression at comparable levels between cell lines (**Additional file 12: Supplementary Figure 5A**). Similarly, *CHAT* expression is only upregulated after differentiation into MNs (**Additional file 12: Supplementary Figure 5A**). Nevertheless, SOD1^D90A-KI^ showed significant downregulation of *CHAT* at 28 days that persisted up to 56 days. Additionally, SOD1^D90A-KI/G93A-KI^ exhibited a significant downregulation of *CHAT* at 56 days. Further analyses at the protein level by immunocytochemistry (**Figure 2A**) did not show statistical differences in the levels of ISL1 (**Figure 2B**), HB9 (**Figure 2C**), or CHAT (**Figure 2D**) between cell lines at 56 days. While slight differences were observed at the transcript level, the expression of MN markers at the protein level within MN spheroids was comparable across cell lines. Together, these findings confirmed that the number of MNs generated within the MN spheroids was comparable between cell lines at 56 days, the end point for these studies.

iPSC differentiation protocols into MNs typically do not yield pure cultures containing only MNs. Previous work has shown the presence of other cell types, including interneurons and glial cells (28, 31, 36). Thus, to investigate the cell identities present in our cultures, we assessed the presence of transcriptional markers associated to interneurons (INs) from the V2 and V3 progenitor domains which flank the MN domain (30), oligodendrocytes and astrocytes (**Additional file 12: Supplementary Figure 5B-E**).

Interneurons (INs) from the V2a domain are identified by the expression of *VSX2* (37). At the iPSC and MNPC stages, *VSX2* expression was nearly absent, and was upregulated at later stages of differentiation. On day 28, there was a trend toward a downregulation of *VSX2* in SOD1^D90A-KI^ MN spheroids, although it did not reach statistical significance. The expression of *NKX2.2*, a marker associated with IN progenitors of the V3 domain, was expressed at the MNPC stage at similar levels between cell lines, and it was subsequently downregulated in MN spheroids at 14 and 28 days. As V3 domain progenitors exited the cell cycle, they differentiated into V3 INs as indicated by *SIM1* expression in MN spheroids. At day 28, although non-significant, there was a trend toward higher levels of *SIM1* in SOD1^D90A-KI^ and SOD1^G93A-KI^ MN spheroids (**Additional file 12: Supplementary Figure 5B**).

Since OLIG2 is also considered a pan-oligodendrocyte gene, we characterized the expression of *OLIG2* (**Supplementary Figure 3**) along with *SOX10* and *MBP* to investigate the presence of oligodendrocyte precursor cells (OPCs) and mature oligodendrocytes (38), respectively. At the iPSC stage, we detected the expression of *SOX10*, however, it was rapidly downregulated to negligible levels at the MNPC and MN stages. In contrast, *MBP* expression reached its peak in MN spheroids at 28 days, with no significant differences observed between the cell lines (**Additional file 12: Supplementary Figure 5C**).

Finally, to assess the presence of astrocytes within our MN spheroids, we checked the expression of established astrocyte markers, *ALDH1L1* (39) and *GFAP* (40). These markers were undetectable in MN spheroids from any cell line at 28 days (**Additional file 12: Supplementary Figure 5D**). Similarly, the pluripotency-associated markers *NANOG* and *OCT4*, which are used to characterize the cells at the iPSC stage (41), were absent in both MNPCs and MN spheroids independent of the cell line (**Additional file 12: Supplementary Figure 5E**).

Altogether, our results confirm the presence of other cellular types within the MN spheroids, namely INs from V2 and V3 domains, and oligodendrocytes. Although some trends were observed, the overall results indicate that there are no significant differences in the proportion of cell types present within MN spheroids across the different cell lines.

### SOD1 expression is altered in SOD1 MN spheroids

As part of the end stage characterization, we assessed the expression of SOD1 in 56-day-old MN spheroids at the transcript and protein level through qPCR, immunostaining, and Western Blot, respectively.

As expected, *SOD1* was expressed at all differentiation stages due to its critical role as ROS scavenger (**Figure 3A**). Given that SOD1 functions in most subcellular compartments (42), it was anticipated that its transcription levels would be higher in differentiated neurons, correlating with the increased cell volume. Only SOD1^G93A-KI^ MN spheroids had a significant upregulation of *SOD1* at 28 days; however, its levels returned to comparable levels with the isogenic control at 56 days. Strikingly, analysis of SOD1 at the protein level revealed distinct expression patterns across cell lines. Immunoblotting against the C-terminal domain of SOD1 showed that SOD1^D90A-KI^ MN spheroids have a strong upregulation of SOD1 (**Figure 3B/C**). In contrast, SOD1^G93A-KI^ and SOD1^D90A/G93A-KI^ MN spheroids displayed a significant downregulation in SOD1 (**Figure 3B/C**). Of note, SOD1^D90A-KI^ and SOD1^D90A/G93A-KI^ monomers displayed a higher motility which is expected given the predicted reduction in molecular mass that results from the Asp-to-Ala change (43, 44). Finally, an immunocytochemistry analysis showed similar results to those observed by Western Blot analysis in 56-day-old MN spheroids (**Figure 3D/E**). Importantly, the antibodies used in this study to detect SOD1 expression by Western Blot and immunocytochemistry were selected based on a recent publication validating numerous commercial SOD1 antibodies with SOD1-KO lines (45).

**Figure 3.**
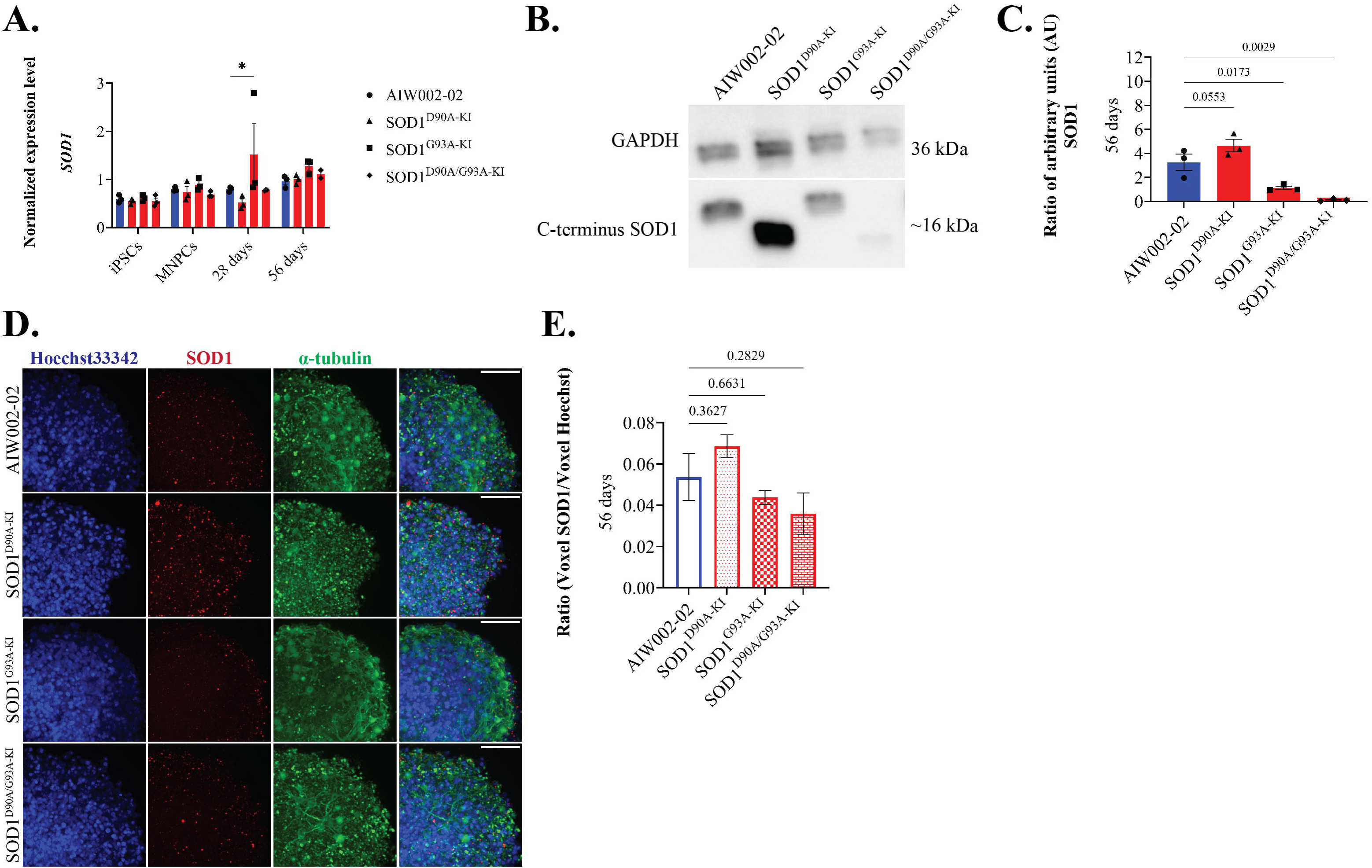
Transcript and protein levels of SOD1 in MN spheroids. **A.** A RT-qPCR was performed to assess the levels of *SOD1* in iPSCs, MNPCs, 28-day-old and 56-day-old MN spheroids. At 56 days in culture, *SOD1* was found at comparable levels in MN spheroids from all lines. Data shown as mean ± SEM, N=3, n=3. Significance was determined using a two-way ANOVA, followed by a post hoc Dunnett’s test using AIW002-02 as the reference sample. *, p ≤ 0.05. **B/C.** Protein quantification of SOD1 through Western Blot analysis showed a strong trend toward upregulation of SOD1 in SOD1^D90A-KI^ MN spheroids. In contrast, SOD1^G93A-KI^ and SOD1^D90A/G93A-KI^ lines showed a mild trend toward downregulation of SOD1. Data shown as mean ± SEM, N=3. **D/E.** Similarly, quantification through immunocytochemistry showed similar SOD1 (red) trends to those observed by Western Blot. Data shown as mean ± SEM, N=3, n=3.

Taken together, our results show that SOD1 expression is dysregulated at the protein level in SOD1 MN spheroids. However, as mutated SOD1 is upregulated in SOD1^D90A-KI^ and downregulated in SOD1^G93A-KI^ and SOD1^D90A/G93A-KI^ MN spheroids, the mechanism through which SOD1 exerts its deleterious effects may differ among cell lines.

### Dysregulation of neurofilament (NF) subunits in SOD1 MN spheroids

Neurofilaments (NF) are composed of three protein subunits that are the most abundant cytoskeletal component of neuronal cells: NF light (NFL), NF medium (NFM), and NF heavy (NFH) (46). Given the growing evidence that NF abnormalities may be an early pathological feature of ALS (47, 48), we characterized the transcriptional expression of the three genes encoding neurofilament subunits—*NEFL*, *NEFM*, and *NEFH*—in iPSCs, MNPCs, and MN spheroids derived from the three SOD1 lines and the isogenic control after 14, 28, and 56 days in culture. As expected, iPSCs from all cell lines lack expression of any NF subunit. Subsequently, MNPCs begin to express NF subunits, albeit in lesser proportions compared to *NES*, which encodes for Nestin, an intermediate filament characteristic of immature neurons (**Additional file 13: Supplementary Figure 5A**). Finally, the expression of NF subunits was increased upon differentiation into MN spheroids.

*NEFL* expression did not show significant differences between cell lines at 28 and 56 days (**Additional file 13: Supplementary Figure 5B**). In contrast, *NEFM* expression showed a significant downregulation in SOD1^D90A-KI^ and SOD1^G93A-KI^ MN spheroids at 28 days (**Additional file 13: Supplementary Figure 5C**). Interestingly, the levels of *NEFM* in SOD1^D90A-KI^ MN spheroids were comparable to the isogenic control at 56 days, while those of SOD1^G93A-KI^ MN spheroids became significantly upregulated. As well, *NEFH* expression was significantly downregulated in SOD1^D90A-KI^ and SOD1^D90A/G93A-KI^ MN spheroids at 28 days (**Additional file 13: Supplementary Figure 5D**). At 56 days, *NEFH* was downregulated in all SOD1 mutants.

To assess if the differences in expression at the transcriptional level were also present at the protein level, we performed Western Blots on total protein extractions of 56-day-old MN spheroids (**Figure 4A**). To perform this assay, we pooled multiple 56-day-old MN spheroids per sample. We observed the downregulation of NFL across all SOD1 mutants, with SOD1^D90A-KI^ MN spheroids reaching statistical significance (**Figure 4B**). Among the three NF protein subunits, NFM exhibited the least variation, showing no significant differences between cell lines (**Figure 4C**). Conversely, NFH displayed a comparable pattern to NFL in SOD1^D90A-KI^ MN and SOD1^D90A/G93A-KI^ MN spheroids, also reaching statistical significance in SOD1^D90A-KI^ MN spheroids (**Figure 4D**). To further investigate if the observed changes were a result of a generalized cytoskeleton disruption, we quantified the expression of βIII-tubulin but no significant changes were found between cell lines (**Figure 4E**).

**Figure 4.**
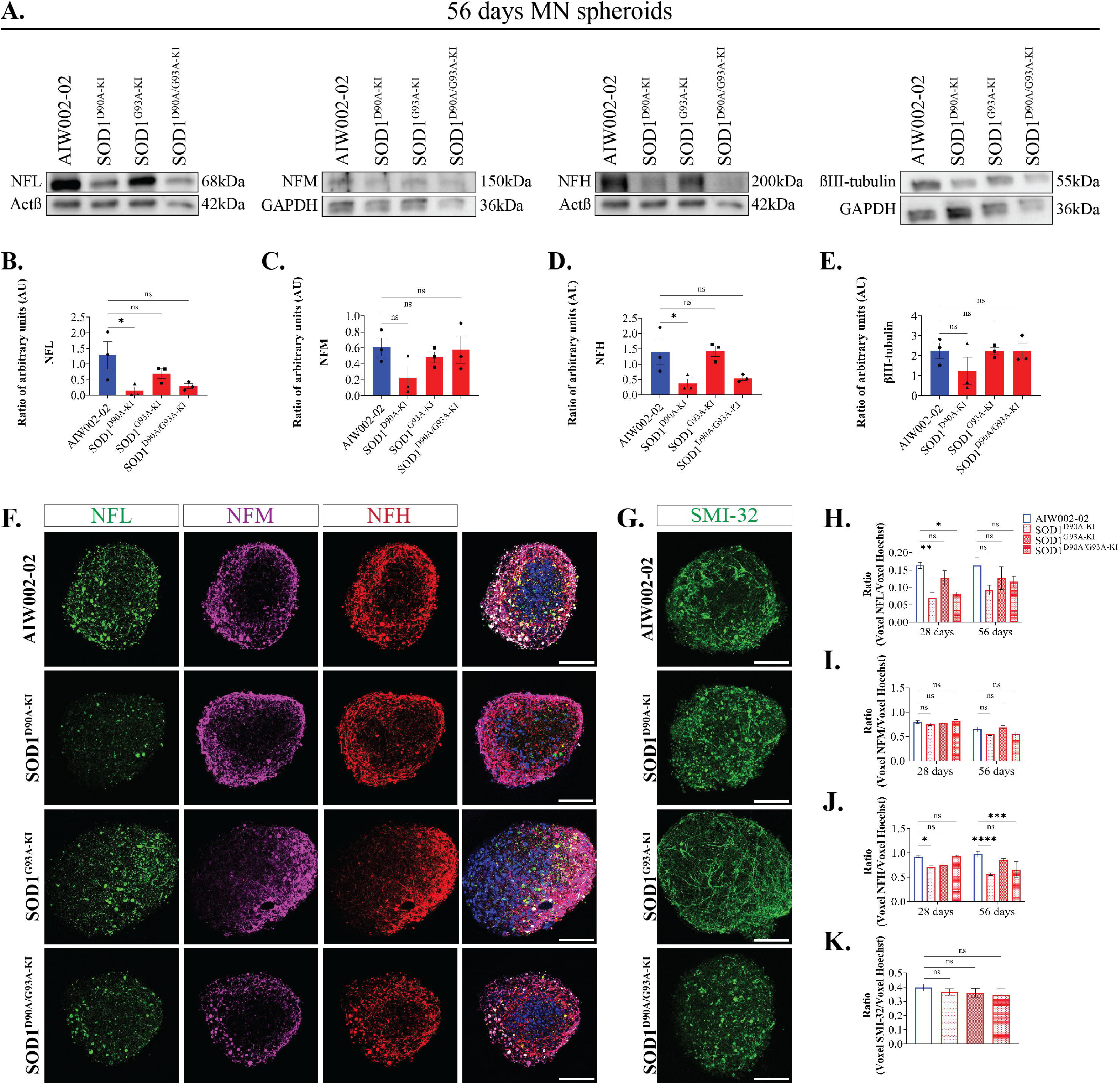
SOD1 MN spheroids showed reduced protein expression of neurofilament (NF) subunits. Western Blot and immunocytochemistry analyses were performed to assess the expression of NF subunits at the protein level. **A.** For Western Blot, sixty spheroids were pooled per sample to quantify the levels of NFL, NFM, NFH, and βIII-tubulin **B.** NFL showed a trend toward downregulation in all SOD1 mutants, with a statistically significant reduction observed in SOD1^D90A-KI^ MN spheroids. **C.** In contrast, NEFM expression remained unchanged across cell lines. **D.** Finally, NEFH exhibited a trend similar to NFL in both SOD1^D90A-KI^ and SOD1^D90A/G93A-KI^ MN spheroids, though significance was reached only in the SOD1^D90A-KI^ line. **E.** βIII-tubulin was used as a control for generalized cytoplasm alterations. Data shown as mean ± SEM, N=3, n=3. Full-length blots are presented in **Additional file 15:** Supplementary Figure 8. **F.** Immunocytochemistry was performed on individual MN spheroids using antibodies against NFL (green), NFM (magenta), NFH (red), and Hoechst33342 (blue). Images represent a single slice from the middle of the Z-stack, acquired through confocal imaging. Scale bar, 100 µm. **G.** A second immunostaining against SMI-32 was conducted to control for the number of MN within MN spheroids. Images represent the maximal projection of 12 optical slices acquired through confocal imaging. Scale bar, 100 µm. **H-K.** Quantification of the immunostainings revealed trends consistent with those observed in Western Blot analyses, where NFL and NEFH were primarily downregulated in SOD1^D90A-KI^ and SOD1^D90A/G93A-KI^ MN spheroids. In contrast, NFM and SMI-32 levels remained stable across all cell lines. Data shown as mean ± SEM, N=3, n=4. Significance was determined using a two-way ANOVA, followed by a post hoc Dunnett’s test using AIW002-02 as the reference sample. *, p ≤ 0.05; **, p ≤ 0.01; ***, p ≤ 0.001; ****, p ≤ 0.0001; ns, non-significant (p > 0.05).

To investigate NF subunit protein expression within individual MN spheroids, we conducted immunocytochemistry analyses in 56-day-old MN spheroids (**Figure 4F**). Additionally, we immunostained the MN spheroids against SMI-32, which targets non-phosphorylated NFH enriched in MNs (**Figure 4G**). As expected, the protein expression levels of NF subunits, as measured by immunocytochemistry, mirrored the trends observed in the Western Blot analyses. NFL was downregulated in MN spheroids from all SOD1 mutants (**Figure 4H**), with statistically significant reductions observed in SOD1^D90A/G93A-KI^ MN spheroids at 28 days and in SOD1^D90A-KI^ MN spheroids at 28 and 56 days. NFM remained the least affected NF subunit, maintaining comparable levels across all cell lines (**Figure 4I**). Lastly, NEFH showed significant downregulation in SOD1^D90A-KI^ MN spheroids at 28 and 56 days, as well as in SOD1^D90A/G93A-KI^ MN spheroids at 56 days (**Figure 4J**). SMI-32 levels remained consistent across cell lines at 56 days (**Figure 4K**), indicating a similar number of MNs within the MN spheroids. Collectively, these findings suggest that SOD1 mutations are associated with dysregulation of NF subunits, which could lead to NF aggregation and neuronal degeneration as previously reported by another group (49).

### SOD1 MN spheroids exhibited reduced viability and neuronal degeneration at 56 days

We next investigated whether the different SOD1 mutations lead to a decrease in viability (including metabolic activity and survival), and/or an increase in neuronal degeneration in 56-day-old MN spheroids.

To assess cell viability, a CellTiter-Glo® 3D cell viability assay and a Western blot for the apoptosis marker cleaved caspase 3 (CC3) were performed. The CellTiter-Glo® 3D cell viability assay (**Figure 5A**) showed a decrease in ATP levels for the three SOD1 cell lines (SOD1^D90A-KI^, 63.5 ± 6.84 %; SOD1^G93A-KI^, 83.31 ± 8.16 %; SOD1^D90A-KI/G93A-KI^, 77.56 ± 7.31 %). However, immunoblotting for CC3 (**Figure 5B/C**), revealed a trend toward increased levels of CC3 in SOD1^G93A-KI^ and SOD1^D90A-KI/G93A-KI^ MN spheroids.

**Figure 5.**
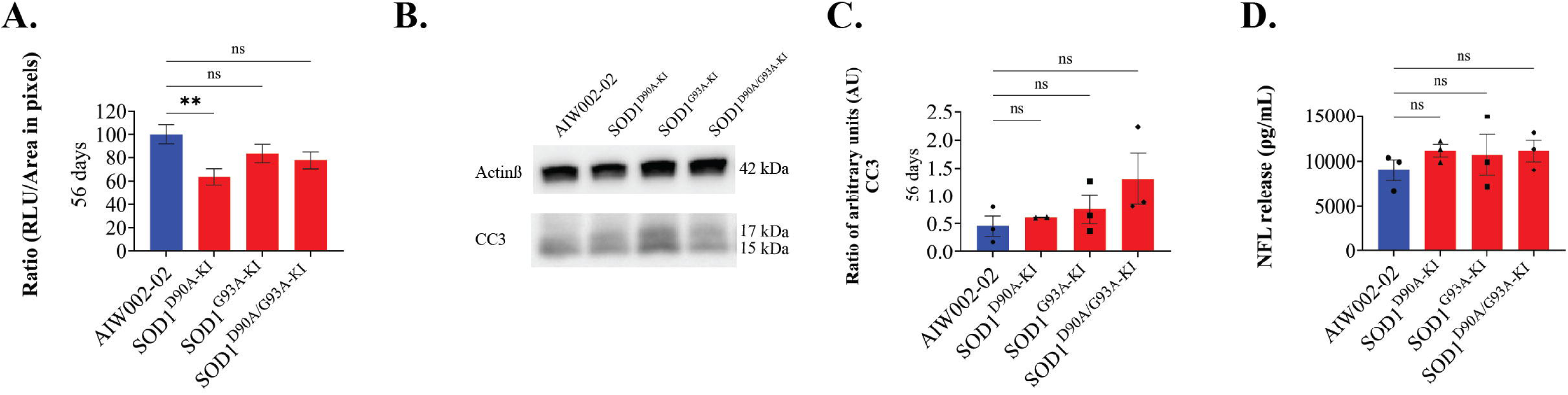
Assessment of cell death and neuronal degeneration indicators in SOD1 MN spheroids. **A.** A CellTiter-Glo® 3D cell viability assay performed in 56-day-old MN spheroids indicated a trend toward decreased cell viability in SOD1 lines with SOD1^D90A-KI^ line reaching statistical significance. Data shown as mean ± SEM, N=3, n=2. **B/C.** Quantification of cleaved-caspase-3 through Western Blot indicated a trend toward increased apoptosis in SOD1^G93A-KI^ and SOD1^D90A/G93A-KI^ lines. Data shown as mean ± SEM, N=3, n=2/3. Full-length blot is presented in **Additional file 15:** Supplementary Figure 8. **D.** Quantification of NFL protein released in the culture media showed a trend toward increased neuronal degeneration in SOD1 lines. Data shown as mean ± SEM, N=3, n=2. Significance was determined using a one-way ANOVA, followed by a post hoc Dunnett’s test using AIW002-02 as the reference sample. **, p ≤ 0.01. ns, non-significant.

In ALS, MN death results in the release of NFL protein into the CSF. Currently, the amount of NFL protein released into CSF, serum, or plasma is often used as a fluid biomarker to indicate disease severity and progression (50, 51, 52). Hence, to further assess the occurrence of neuronal degeneration in our model, we conducted an ELISA assay to measure NFL release in media that had been in contact with the MN spheroids for 56 days. To validate the ELISA assay, we assessed NFL release in the plasma samples from two healthy donors (one female and one male) and two sporadic ALS patients (one female and one male). The ages of the donors ranged from 55 to 70 years (**Additional file 14: Supplementary Figure 7A**). Consistent with findings in ALS patients, extracellular NFL levels were higher in the plasma of sporadic ALS patients compared to control donors (**Additional file 14: Supplementary Figure 7B**) in our assay. Extending this assay into MN spheroids, we observed an increase in NFL levels in the media for all three SOD1 cell lines (AIW002-02, 9000 ± 1144 pg/mL; SOD1^D90A-KI^, 11121 ± 660 pg/mL; SOD1^G93A-KI^, 10667 ± 2263 pg/mL; SOD1^D90A/G93A-KI^, 11121 ± 1183 pg/mL) (**Figure 5D**).

The findings across all three assays indicate that SOD1 mutations are linked to decreased cell viability and increased neuronal degeneration in iPSC-derived MN spheroids.

### Hypoactivity and irregular burst activity are linked to SOD1 mutations

Disrupted neuronal activity has been linked to ALS using iPSC-derived MNs (18, 53, 54). Given that our MN spheroids exhibited reduced viability, increased neuronal degeneration, and downregulation of NF subunits—recently identified as integral components of synapses—we sought to evaluate the functional activity of our MN spheroids using an MEA approach. For this, 7 day old MN spheroids were plated onto electrodes into MEA plates to perform longitudinal recordings until day 56 post-plating (p-p) (**Figure 6A**). The mean firing rate (MFR), representing the average number of spikes per second (Hz), decreased in all SOD1 mutants from 28 days p-p onward. Notably, SOD1^G93A-KI^ MN spheroids were significantly affected after 42 and 56 days p-p (**Figure 6B**).

**Figure 6.**
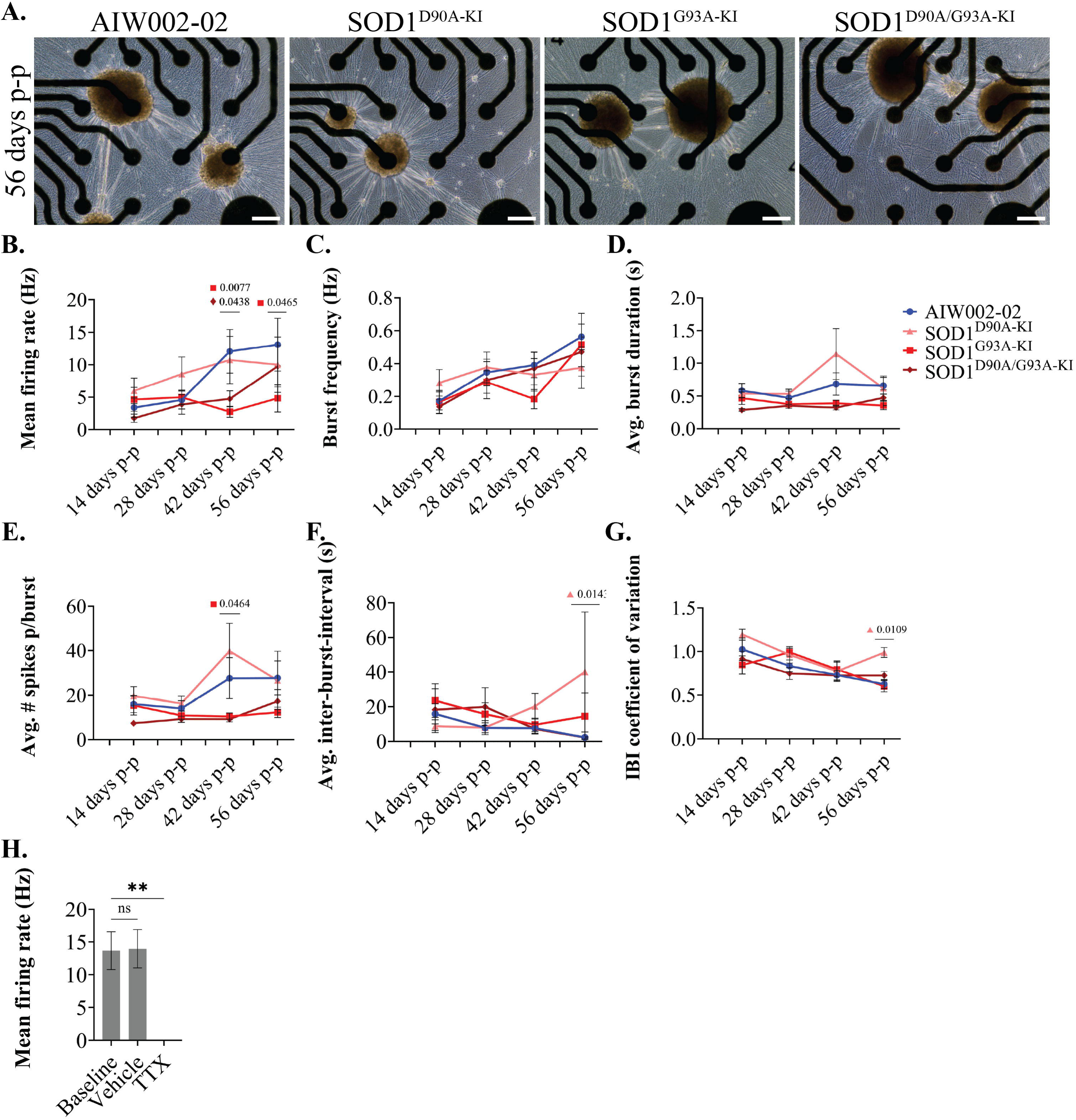
MEA recordings revealed hypoactivity and irregular burst properties in SOD1 MN spheroids. **A.** Representative bright-field images of iPSC-derived MN spheroids at 56 days p-p in Cytoview 24–well MEA plates. Scale bar, 200 µm. **B**. The MFR (Hz) was significantly reduced in SOD1^D90A/G93A-KI^ MN spheroids at 42 days p-p, and in SOD1^G93A-KI^ MN spheroids at both 42 and 56 days p-p. **C.** Burst frequency (Hz) showed a slight reduction in SOD1 MN spheroids at 56 days p-p, although no statistically significant differences were observed. **D.** Starting at 28 days, SOD1^G93A-KI^ and SOD1^D90A/G93A-KI^ MN spheroids displayed reduced average burst duration. **E.** Also, a reduced number of spikes per burst was recorded in SOD1^G93A-KI^ and SOD1^D90A/G93A-KI^ MN spheroids from 28 days onward. **F**. Consistent with the reduced number of bursts observed in SOD1 MN spheroids, the SOD1^D90A-KI^ line showed a significant increase in the average inter-burst interval at 56 days. **G.** Additionally, the IBI coefficient of variation was significantly elevated in SOD1^D90A-KI^ MN spheroids a 56 days, indicating irregular burst activity compared to the isogenic control. Data shown as mean ± SEM, N=3, n=3–10. Significance was determined using a two-way ANOVA, followed by a post hoc Dunnett’s test using AIW002-02 as the reference sample. **H.** Acute treatment with vehicle or TTX was administered to all cell lines to verify the absence of recording artifacts. Data pooled from all cell lines and shown as mean ± SEM, n=3–9. Significance was determined using a one-way ANOVA, followed by a post hoc Dunnett’s test using Baseline as the reference sample. *, p ≤ 0.05; **, p ≤ 0.01; ns, non-significant (p > 0.05).

Next, we assessed burst frequency per MN spheroid and various detailed properties of burst activity (**Figure 6C-G**). Burst frequency, defined as the average number of bursts per second (Hz), showed a slight reduction in SOD1 MN spheroids at 56 days p-p; however, no statistically significant differences were detected (**Figure 6C**). Starting at 28 days, SOD1^G93A-KI^ and SOD1^D90A/G93A-KI^ MN spheroids showed reduced average burst duration (**Figure 6D**) and a lower average number of spikes per burst (**Figure 6E**). Considering the reduced number of bursts observed in SOD1 MN spheroids, an increase in the average inter-burst-interval (s) was anticipated. This increase reached statistical significance for SOD1^D90A-KI^ at 56 days, whereas no significant changes were observed for SOD1^G93A-KI^ and SOD1^D90A/G93A-KI^ MN spheroids (**Figure 6F**). The inter-burst interval (IBI) coefficient of variation quantifies the variability in burst timing. An IBI coefficient of variation near zero indicates regular, consistent bursting, while higher values typically reflect irregular or disrupted neuronal activity. Throughout our longitudinal recordings, the IBI coefficient of variation was significantly elevated in SOD1^D90A-KI^ MN spheroids at 56 days, indicating irregular burst activity compared to the isogenic control (**Figure 6G**). Importantly, acute treatment with tetrodotoxin (TTX) completely abolished the activity of MN spheroids across cell lines, confirming that our observations were not artifacts of the recording system (**Figure 6H**).

Overall, we observed a phenotype of hypoactivity in MN spheroids with SOD1 mutations, becoming apparent after 28 days p-p, with SOD1^G93A-KI^ MN spheroids being the most severely affected. Interestingly, although SOD1^D90A-KI^ MN spheroids did not show the most pronounced changes in overall activity compared to the isogenic control, they exhibited significant alterations in burst properties. Collectively, our findings suggest that SOD1 mutations are associated with hypoactivity and irregular burst activity.

## Discussion

In this study, we used a protocol previously optimized by our group (23) to generate and characterize iPSC-derived MN spheroids from iPSC lines harboring homozygous mutations in *SOD1* and their corresponding isogenic control (AIW002-02). Notably, our model is highly scalable, allowing for the processing and analysis of multiple spheroids simultaneously, while also supporting long-term cultures for up to 56 days.

First, we performed the size profiling of our MN spheroids to assess potential morphological changes. SOD1^D90A-KI^ MN spheroids were smaller, while SOD1^G93A-KI^ and SOD1^D90A/G93A-KI^ were larger relative to the isogenic control. We hypothesized that size disparities arise from differences in the rate at which neural precursors exit the mitotic phase and commit to differentiation into neurons. To investigate this, we analyzed the downregulation of SOX1, a marker of pan-neuronal proliferative precursors. Surprisingly, SOD1^D90A-KI^ MN spheroids exhibited the highest expression of SOX-1, despite their smaller size, while SOD1^G93A-KI^ MN spheroids showed the lowest expression of SOX-1, regardless of their larger size. Only in the case of SOD1^D90A/G93A-KI^ MN spheroids could we conclude that the larger size was attributable to a slower rate of SOX1 downregulation. Although this result is intriguing, understanding the mechanisms driving the subtle size differences in MN spheroids falls outside the scope of this work. It is important to note, however, that the overall morphology of the MN spheroids and the expression of MN markers remain consistent across cell lines.

Studies using iPSC-derived MNs as a model have reported either increased or unchanged SOD1 expression, depending on the specific mutation analyzed (49, 53, 55). Here, we observed an upward trend toward higher levels of SOD1 in SOD1^D90A-KI^ MN spheroids, although no statistical differences were detected. Similarly, the assessment of SOD1 expression in post-mortem tissue of the central nervous system and peripheral organs of ALS patients with a homozygous D90A mutation did not report significant differences compared to controls (43). In contrast, the use of reliable antibodies for SOD1 detection (45) allowed us to report significantly reduced SOD1 levels in SOD1^G93A-KI^ and SOD1^D90A/G93A-KI^ MN spheroids. A separate study on iPSC-derived MNs with a heterozygous G93A knock-in mutation reported unchanged SOD1 levels in the RIPA-soluble fraction compared to the isogenic control (53), prompting the question of whether differences in zygosity explain the conflicting results. Overall, ALS patients with SOD1 mutations have significantly reduced SOD1 protein levels, with varying degrees of protein expression changes associated with different SOD1 variants (10). Additionally, it is worth noting that the large validation study conducted by Ayoubi et al. (2023), enabled us to use reliable antibodies for SOD1 detection (45). Notably, the study conducted by Ayoubi et al. (2023) revealed that several SOD1 antibodies produce false-positive results, potentially confounding earlier findings, and underscores the value of Open Science initiatives that conduct such validations for the scientific community (45).

Cell viability and neuronal degeneration are commonly used as indicators of ALS pathology in iPSC-derived cells. In this study, we performed three assays to assess these indicators within 56-day-old MN spheroids. Initially, a 3D CellTiter-Glo assay was performed to measure ATP levels in MN spheroids, as higher ATP levels indicate active metabolism and viable cells. We observed reduced ATP levels in all SOD1 lines compared to the isogenic control, confirming that SOD1 mutations lead to a reduction in metabolic activity, thereby impacting cell viability. Next, we conducted a Western blot analysis for cleaved caspase-3, a downstream protein fragment in the apoptotic pathway that has been previously reported to be upregulated in iPSC-derived motor neurons with SOD1 mutations (56). As anticipated, we observed upregulation of □15 and □17 kDa protein fragments but only in SOD1^G93A-KI^ and SOD1^D90A/G93A-KI^ MN spheroids. Surprisingly, SOD1^D90A-KI^ MN spheroids showed cleaved caspase-3 levels comparable to the isogenic control, despite exhibiting the lowest ATP levels in the CellTiter-Glo assay. A plausible explanation is that, while SOD1^D90A-KI^ MN spheroids are presented with reduced metabolic activity, they have not yet entered apoptosis. Cell viability remains a debated topic in ALS studies using iPSC-derived motor neurons, with some studies reporting a decrease in cell viability and activation of the apoptotic pathway (56, 57, 58), while others find no significant change (18, 59). It has been suggested that the failure to observe changes in viability is due to the technical challenge of maintaining 2D cultures in optimal conditions for long-term analysis, an important factor for studying age-associated progressive diseases (18, 60). Our study confirms that using a 3D culture system can sustain cells for longer periods, thereby revealing delayed phenotypes.

To assess neuronal degeneration, we investigated NFL subunit release into the media of MN spheroids, as elevated levels of NFL in blood, serum, plasma, and CSF are established biomarkers of ALS, indicative of axonal damage associated with the disease (51, 61). This approach has undergone limited optimization for cultured cells, particularly iPSC-derived cells, which presented a challenge for this assay. Therefore, to validate our method, NFL levels were also measured in plasma samples from healthy donors and ALS patients. All SOD1 lines trended toward an increase in NFL subunit levels in the media, although this was outside of statistical significance. We attribute this to NFL subunit release being measured from the pooled media of six MN spheroids, each composed of 10K cells, which may produce NFL levels that are only marginally above the detection limit of the measuring kit. Increasing the size of the MN spheroids, pooling media from additional samples and/or increasing the culture time can potentially lead to a significant increase in NFL release, given how ALS is a progressive disease.

Studies investigating NF subunit expression in post-mortem tissues from sporadic ALS patients (62, 63) and iPSC-derived MNs with SOD1 mutations (49) have reported reduced NFL expression at both transcript and protein levels. Supporting these findings, evidence indicates that mutant SOD1 directly binds to *NEFL* mRNA, disrupting its stability (49, 64). In contrast, inconsistent results were reported when analyzing the expression of NFM and NFH in diseased MNs. While NFM and NFH protein expressions are downregulated in SOD1 iPSC-derived MNs (49), they are either upregulated at the protein level (65) or unaffected at the transcriptional level (63) in post-mortem tissue from sporadic ALS patients. More recently, a study analyzing the transcriptome in post-mortem spinal cord tissue of SOD1 ALS patients via bulk RNA-seq revealed significant downregulation of *NEFL*, *NEFM*, and *NEFH* (*66*). In our study, SOD1^D90A-KI^ MN spheroids exhibited significant reductions in NFL at the protein level, in *NEFM* at the transcriptional level, and in NFH at both the transcriptional and protein levels. Similarly, SOD1^D90A/G93A-KI^ MN spheroids showed a trend toward downregulation of NFL and a significant reduction in NFH at the protein level. Interestingly, SOD1^G93A-KI^ MN spheroids were the least affected, showing only a trend toward downregulation of NFL at the protein level. Overall, NFL appears to be the most downregulated NF subunit, showing consistent patterns across all SOD1 lines. Taken together, our results align with those reported by Chen et al. (2014), who observed a downregulation of NFL, NFM, and NFH at the protein level in iPSC-derived MNs carrying the A4V and D90A mutations in SOD1. We found it surprising that SOD1^D90A/G93A-KI^ MN spheroids did not exhibit the lowest levels of NF subunits, as we had anticipated a synergistic effect from the presence of both mutations. It appears that the G93A mutation may lead to a distinct interaction between mutated SOD1 and NF subunits, as this cell line was the least affected. Future studies should examine the impact of a broader range of SOD1 mutations on NF subunit regulation.

Finally, as previous studies with iPSC-derived MNs suggest that synaptic dysfunction may precede MN degeneration (18, 59), we conducted a functional analysis using MEA recordings. For all functional properties measured, differences between cell lines emerged after 28 days p-p, emphasizing the importance of using a model that supports long-term culture for phenotypic comparisons. SOD1^G93A-KI^ MN and SOD1^D90A/G93A-KI^ MN spheroids exhibited a hypoactive phenotype while SOD1^D90A-KI^ MN spheroids displayed notable changes in burst properties. Similarly, previous work reported hypoactivity and disrupted burst activity in 2D iPSC-derived MNs with heterozygous SOD1 G93A and A4V mutations when seeded on a monolayer of primary astrocytes (53). In contrast, another study reported hyperexcitability in 2D iPSC-derived MNs carrying the heterozygous SOD1 A4V mutation (54); however, in this study, MNs were differentiated and recorded as a pure 2D culture. In our model, we cannot exclude the possibility that the 3D environment may alter the functional profile of the MNs. Additionally, while INs and oligodendrocytes are present in similar proportions across cell lines, their potential role in modulating MN activity cannot be disregarded as growing evidence suggests their involvement in ALS (67, 68).

The D90A and G93A mutations, both linked to fALS, occur near the β-barrel motif of SOD1 (7). Notably, these mutations are considered WT-like fALS mutations as they maintain crucial structural interactions required for metal-induced stabilization (69). Given these characteristics, we hypothesized that single homozygous mutations would induce similar pathological alterations, with the double mutation possibly exhibiting synergistic effects. Generating a model with enhanced deleterious phenotypes could help to provide deeper insights into the mechanisms of disease. However, only a reduction in cell viability and downregulation of NFL were consistently observed across all three phenotypes, and these changes were not worsened by the double SOD1 mutation. This suggests that the conformational change created by the two mutations may have resulted in a structure similar to that of either of the single SOD1 mutants.

One of the primary challenges in ALS research is its intrinsic heterogeneity. Mutated SOD1 demonstrates variable behavior depending on the specific mutation (11, 12), and patients with fALS caused by SOD1 mutations exhibit diverse clinical manifestations and prognoses (10). This highlights the necessity of analyzing ALS at the genetic level, as even mutations in close proximity or with conserved structural interactions can result in distinct disease mechanisms.

## Conclusion

In summary, we developed a 3D iPSC-derived MN spheroid model that recapitulates key ALS pathology features when SOD1 mutations are present. Our results revealed consistent reductions in cell viability and NFL downregulation across all phenotypes. While MN activity was also affected, hypoactivity was observed in SOD1 G93A and D90A/G93A genotypes, whereas a disrupted bursting pattern was more characteristic of the SOD1 D90A genotype. Notably, for the first time, we demonstrated that two SOD1 mutations linked to disease onset do not exhibit synergistic effects. These findings provide valuable insights into mutation-specific phenotypes and help contribute to our understanding of ALS heterogeneity.

## Declarations

### Ethics approval

The use of iPSCs in this research was approved by the McGill Research Ethics Board (IRB Study Number A03-M19-22A).

### Consent for publication

Not applicable.

### Availability of data and materials

All data generated or analyzed during this study are included in this published article.

### Authors’ contributions

MJC-M contributed to the conceptualization, study design, data collection, result analysis, and the writing, reviewing, and editing of the manuscript. AKF-F contributed significantly to data collection, result analysis, and the reviewing and editing of the manuscript. MN generated and validated the CRISPR/Cas9-edited iPSC lines. GH generated and characterized iPSC-derived MNPCs. SL developed image analysis pipelines. MB-T contributed with image analysis. CX-QC and TMG performed quality control analyses for all iPSC lines. NA performed the karyotyping of all iPSC lines. DH contributed to image acquisition and analysis. MC contributed to the analysis of results, funding acquisition, as well as the reviewing and editing of the manuscript. LG generated and characterized AIW002-02 iPSC-derived MNPCs. NA conducted quality control analyses for the AIW002-02 iPSC line. ED participated in generating the CRISPR/Cas9-edited iPSC lines. TMD contributed to the study design, funding acquisition, and the reviewing and editing of the manuscript. All authors have read and agreed to the published version of the manuscript.

## Acknowledgements

We thank the donors for their generous donations of samples. Their contributions are invaluable in advancing our understanding of ALS. We thank The Neuro’s Open Biobank (or Clinical Biospecimen Imaging and Genetic Repository, C-BIG), directed by Dr. Karamchandani, for providing us with access to the plasma samples from control and patient donors used for this study. We thank YCharOS for their expertise in antibody validation and the donation of different SOD1 antibodies for troubleshooting. We thank Martin H. Berryer and Mark Aurousseau for their valuable input in the drafting of this manuscript.

