## Additional File 1 for "Mutations in SOD1 induce ALS-related phenotypes in 3D iPSC-derived motor neuron (MN) spheroids"

**CRISPR/Cas9 genome editing and validation**

Following a streamlined CRISPR workflow (1, 2), the P3 Primary Cell-4D Nucleofector™ X Kit S (#V4XP-3032, Lonza) was used to correct either p.D90A or p.G93A in the SOD1^D90A/G93A-KI^ iPSC line, thereby generating two iPSC lines with a single *SOD1* mutation. The sequences of the sgRNAs and ssODNs used are provided in **Table S2**.

After isolation, gene-edited clones were identified by ddPCR with a QX200™ Droplet Reader (Bio-Rad). The detection of the modified nucleotides by ddPCR was based on a TaqMan® assay including two PCR primers: one DNA probe fused to a fluorophore specific to the corrected/WT allele (FAM), and one DNA probe fused to a fluorophore specific to the edited allele (HEX). Locked Nucleic Acid (LNA®) probes were designed following the manufacturer’s criteria. Sanger sequencing was used to validate the sequence integrity of successful clones. The sequences of the primers and probes used for ddPCR, and Sanger sequencing are provided in **Table S3**.

**Karyotyping**

For G-band karyotyping, AIW002-02 and SOD1^D90A/G93A-KI^ iPSCs were cultured for 72 h until they reached 50–60% confluency. iPSCs were split, pellet, and shipped live to the Wicell Cytogenetics Core (instructions provided by WiCell, Madison, WI, USA).

SOD1^D90A-KI^ and SOD1^G93A-KI^ iPSCs were cultured for 72 h until they reached 50–60 % confluency, and karyotyping was conducted as previously described (3).

**Genomic abnormalities analyses**

Genomic DNA was extracted with the Genomic DNA Mini Kit (Geneaid) according to the manufacturer's instructions. Genomic stability was detected using the hPSC Genetic Analysis Kit (#07550, STEMCELL Technologies). Briefly, 5 ng of genomic DNA was mixed with a ROX reference dye, and double-quenched probes tagged with 5-FAM that represent the eight most common karyotypic abnormalities reported in hiPSC: chr 1q, chr 8q, chr 10p, chr 12p, chr 17q, chr 18q, chr 20q or chr Xp. Reactions were run on a QuantStudio 5 Real-Time PCR System (Thermo Fisher Scientific). Copy numbers were analyzed using the ΔΔCT method and the copy number of a control region in chr4 was used for normalization (4, 5).

**Virology test**

Blood samples were cleared from Hepatitis A/B and HIV at the clinic. After reprogramming, iPSCs are assessed for mycoplasma using the MycoAlert® Mycoplasma Detection Kit (#LT07-318, Lonza) following manufacturer’s instructions.

**Immunocytochemistry of cell monolayers grown on glass coverslips**

Immunocytochemistry of cell monolayers (i.e. iPSCs and MNPCs) grown on glass coverslips (#41001112, Fisher Scientific) was performed as previously described (4). Briefly, cells were fixed in 4% FA diluted in 1X PBS for 15–20 min at room temperature (RT) and washed three times for 5 min with 1X PBS. Cells were permeabilized with 0.2% Triton X-100 (#TRX506, Bioshop) diluted in 1X PBS for 10 min at RT, and were then blocked for 1 h at RT in a blocking solution containing 5% normal donkey serum (NDS; #S30 Normal donkey, Millipore), 1% bovine serum albumin (BSA; #800-095-CG, Wisent Bioproducts), and 0.05% Triton X-100 diluted in 1X PBS (blocking solution). After blocking, cells were incubated with primary antibodies diluted in blocking solution overnight at 4°C. Primary antibodies were washed out by performing three 5 min washes with 1X PBS and cells were subsequently incubated with secondary antibodies diluted in blocking solution for 2 h at RT. Secondary antibodies were washed out by performing three 5 min washes with 1X PBS followed by a Hoechst33342 nucleic acid counterstain for 5 min. Coverslips were mounted with Fluoromount-G^TM^ (00-4958-02, Thermo Fisher Scientific). Immunocytochemistry images were acquired using the automated Evos FL-Auto2 imaging system (Thermo Fisher Scientific) using 20X magnification (N.A 0.4) or the Zeiss Axio Observer Z1 Inverted Microscope using 20X magnification (N.A 0.8).

**MNPC quantification**

For each cell line, three images per batch were acquired with a Zeiss Axio Observer Z1 Inverted Microscope using 20X magnification (N.A 0.8). A CellProfiler (version 4.2.5) pipeline was used to quantify the number of Olig2^+^, Pax6^+^, and Olig2^+^/Pax6^+^ cells. Briefly, nuclei immunolabelled with Hoechst33342 were identified as primary objects called “Nuclei” based on the global threshold strategy, the Otsu thresholding method, and the diameter in pixels. The pixel intensity within each primary object was measured in the Pax6 channel and a threshold was applied to determine Pax6^+^ cells. The newly generated objects were called “Pax6positive”. A similar approach was used to identify Olig2^+^ cells using the Olig2 channel, and the generated objects were called “Olig2positive”. Finally, the pixel intensity of each “Pax6positive” object was measured in the Olig2 channel, and a threshold was applied to identify double-positive cells. AIW002-02: B1 (125 cells) , B2 (400 cells), B3 (237 cells); SOD1^D90A-KI^: B1 (487 cells), B2 (352 cells), B3 (484 cells); SOD1^G93A-KI^; B1 (209 cells), B2 (292 cells), B3 (405 cells); SOD1^D90A/G93A-KI^: B1 (129 cells), B2 (267 cells), B3 (155 cells).
