## Additional File 2-Supplementary Table 1 for "Mutations in SOD1 induce ALS-related phenotypes in 3D iPSC-derived motor neuron (MN) spheroids"

**Supplementary Table 1. Overview of iPSC lines**

| Cell line ID | ALS mutation | Sex | Age | Ethnicity | Primary cell line | Reprogramming method | Ref. |
| --- | --- | --- | --- | --- | --- | --- | --- |
| AIW002-02 | None | Male | 37 | Caucasian | PBMCs | Sendai virus | (1) |
| *SOD1*^D90A/G93A-KI^ | p. D90A and p. G93A | Male | 37 | Caucasian | AIW002-02 | n/a | (2) |
| *SOD1*^D90A-KI^ | p. D90A | Male | 37 | Caucasian | *SOD1*^D90A/G93A^**^-^**^KI^, G93A correction | n/a |  |
| *SOD1*^G93A-KI^ | p. G93A | Male | 37 | Caucasian | *SOD1*^D90A/G93A^**^-^**^KI^, D90A correction | n/a |  |

1.	Chen CX, Abdian N, Maussion G, Thomas RA, Demirova I, Cai E, Tabatabaei M, Beitel LK, Karamchandani J, Fon EA, Durcan TM. A Multistep Workflow to Evaluate Newly Generated iPSCs and Their Ability to Generate Different Cell Types. *Methods Protoc*. **2021**;4(3).

2.	Deneault E, Chaineau M, Nicouleau M, Castellanos Montiel MJ, Franco Flores AK, Haghi G, Chen CX, Abdian N, Shlaifer I, Beitel LK, Durcan TM. A streamlined CRISPR workflow to introduce mutations and generate isogenic iPSCs for modeling amyotrophic lateral sclerosis. *Methods*. **2022**;203:297-310.
