## Additional File 3-Supplementary Table 2 for "Mutations in SOD1 induce ALS-related phenotypes in 3D iPSC-derived motor neuron (MN) spheroids"

**Supplementary Table 2**. **List of sgRNAs and ssODNS used to make *SOD1* single mutation lines**

| Cell line ID | gRNAs | ssODN template |
| --- | --- | --- |
| *SOD1*^D90A-KI^ | CACAUCGGCCACAGCAUCUU | ATCTGATGCTTTTTCATTATTAGGCATGTTGGAGACTTGGGCAATGTGACTGCTGCCAAAGATGGTGTGGCCGATGTGTCTATTGAAGATTCTGTGATCTCACTCTCAGGAGAC |
| *SOD1*^G93A-KI^ |  | ATCTGATGCTTTTTCATTATTAGGCATGTTGGAGACTTGGGCAATGTGACTGCTGACAAAGATGCTGTGGCCGATGTGTCTATTGAAGATTCTGTGATCTCACTCTCAGGAGAC |
