## Additional File 4-Supplementary Table 3 for "Mutations in SOD1 induce ALS-related phenotypes in 3D iPSC-derived motor neuron (MN) spheroids"

**Supplementary Table 3. List of primers and affinity probes used for ddPCR or Sanger sequencing**

|  | *SOD1*^D90A-KI^ | *SOD1*^G93A-KI^ |
| --- | --- | --- |
| Probe-HEX (mutant)* | /5HEX/AG+AT+G+C+TG+T+GG/3IABkFQ/ | /5HEX/ACT+GCT+G+C+CAAAGA/3IABkFQ/ |
| Probe-FAM (corrected/WT) * | /56-FAM/AGAT+G+G+TG+T+GG/3IABkFQ/ | /5HEX/ACT+GCT+G+A+CAAA+GA/3IABkFQ/ |
| ddPCR primer – F | TTAGTGGCATCAGCCCTAATC | |
| ddPCR primer – R | AGTGTGCGGCCAATGAT | |
| Sanger sequencing primer – F | TCTGAAATCAGGTGCAGCCC | |
| Sanger sequencing primer - R | ACCGCGACTAACAATCAAAGTG | |

*** “+”** signs in front of nucleotides indicate the location of Locked Nucleic Acids (LNA®).
