## Additional File 5-Supplementary Table 4 for "Mutations in SOD1 induce ALS-related phenotypes in 3D iPSC-derived motor neuron (MN) spheroids"

**Supplementary Table 4. List of primary antibodies**

| Antibody | Host | Company | Clone or Cat. # | Dilution (Application) |
| --- | --- | --- | --- | --- |
| Actin | Mouse | Millipore | MAB1501 | 1:40000 (WB) |
| βIII-tubulin | Mouse | Millipore | MAB5564 | 1:20000 (WB) |
| ChAT | Goat | Millipore | MAB144P | 1:50 (ICC) |
| Cleaved-caspase 3 | Rabbit | Cell signaling | 9661 | 1:500 (WB) |
| GAPDH | Mouse | Proteintech | 60004-1-lg | 1:40000 (WB) |
| Hb9 | Mouse | DSHB | 81.5C10 | 1:50 (ICC) |
| Isl-1 | Mouse | DSHB | 40.2D6 | 1:50 (ICC) |
| Ki67 | Mouse | BD Bioscience | 556003 | 1:200 (ICC) |
| Nanog | Rabbit | Abcam | ab21624 | 1:200 (ICC) |
| NFH | Chicken | Abcam | 4680 | 1:1000 (ICC),  1:5000 (WB) |
| NFL | Mouse | Sigma | N5139 | 1:1000 (ICC),  1:5000 (WB) |
| NFM | Rabbit | Millipore | AB1987 | 1:1000 (ICC),  1:5000 (WB) |
| Oct3/4 | Goat | Santa Cruz | Sc-8628 | 1:500 (ICC) |
| Olig2 | Rabbit | Millipore | AB9610 | 1:100 (ICC) |
| Pax6 | Mouse | DSHB | AB_528427 | 1:100 (ICC) |
| SMI-32 | Mouse | Biolegend | 801701 | 1:100 (ICC) |
| SOD1 | Rabbit | Thermo Fisher Scientific | 702783 | 1:500 (ICC) |
| SOD1 (C-terminal) | Rabbit | GeneTex | GTX100659 | 1:500 (WB) |
| SSEA-4 | Mouse | Santa Cruz | sc-21704 | 1:200 (ICC) |
| Tra-1-60 | Mouse | STEMCELL Technologies | TRA1-60R | 1:200 (ICC) |
