## Additional File 6-Supplementary Table 5 for "Mutations in SOD1 induce ALS-related phenotypes in 3D iPSC-derived motor neuron (MN) spheroids"

**Supplementary Table 5. List of secondary antibodies**

| Antibody | Species Reactivity | Company | Cat. # | Dilution (Application) |
| --- | --- | --- | --- | --- |
| Hoechst33342 | n/a | Invitrogen | H3570 | 1:1000 or 1:5000 (ICC) |
| Dylight488 | Rabbit IgG | Abcam | ab96891 | 1:500 (ICC) |
| Dylight650 | Chicken IgY | Abcam | ab96950 | 1:500 (ICC) |
| Dylight488 | Mouse IgG | Abcam | ab96875 | 1:500 (ICC) |
| AlexaFluor647 | Goat IgG | Invitrogen | A21447 | 1:500 (ICC) |
| Dylight550 | Goat IgG | Abcam | ab96936 | 1:500 (ICC) |
| Dylight650 | Mouse IgG | Abcam | Ab96878 | 1:500 (ICC) |
| HRP | Mouse | Jackosn Immunoresearch | 115-035-003 | 1:10000 (WB) |
| HRP | Rabbit | Jackosn Immunoresearch | 111-035-144 | 1:10000 (WB) |
| HRP | Chicken | Jackosn Immunoresearch | 703-035-155 | 1:10000 (WB) |
