## Additional File 7-Supplementary Table 6 for "Mutations in SOD1 induce ALS-related phenotypes in 3D iPSC-derived motor neuron (MN) spheroids"

**Supplementary Table 6. List of TaqMan qPCR probes**

| GENE | Reference |
| --- | --- |
| *GAPDH* | Hs02786624_g1 |
| *ACTB* | Hs01060665_g1 |
| *HOXA5* | Hs00430330_m1 |
| *HOXB8* | Hs00256885_m1 |
| *FOXG1* | Hs01850784_s1 |
| *NKX6.1* | Hs00232355_m1 |
| *KI67* | Hs010327443_m1 |
| *PAX6* | Hs01088114_m1 |
| *OLIG2* | Hs00377820_m1 |
| *NESTIN* | Hs04187831_g1 |
| *HB9* | Hs00907365_m1 |
| *ISL1* | Hs00158126_m1 |
| *CHAT* | Hs00758143_ml |
| *SOD1* | Hs00533490_m1 |
| *NEFH* | Hs00606024_m1 |
| *NEFM* | Hs00193572_m1 |
| *NEFL* | Hs00196245_m1 |
