## Additional File 8-Supplementary Figure 1 for "Mutations in SOD1 induce ALS-related phenotypes in 3D iPSC-derived motor neuron (MN) spheroids"

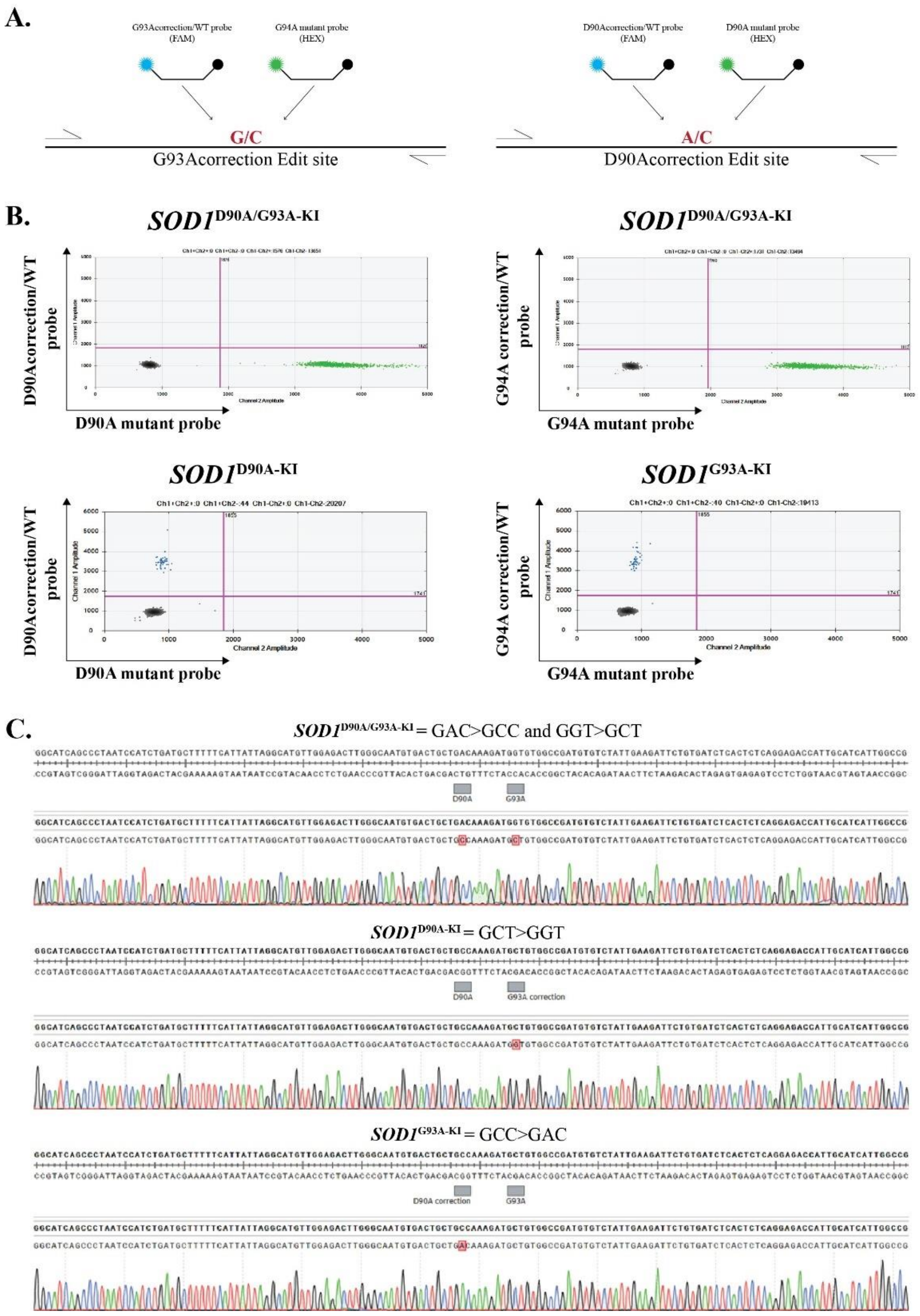


**Supplementary Figure 1. Validation of CRISPR/Cas9 gene editing by ddPCR and Sanger sequencing. A.** Pairs of corrected/WT (FAM, blue) and mutant (HEX, green) probes designed to target the wild-type (WT) or edited alleles, respectively. **B.** ddPCR scatter plots confirming correct gene editing and homozygosity of iPSC lines. **C.** Sanger sequence chromatograms from selected iPSC clones; red box shows complete correction of target nucleotide.
