## Additional File 9-Supplementary Figure 2 for "Mutations in SOD1 induce ALS-related phenotypes in 3D iPSC-derived motor neuron (MN) spheroids"

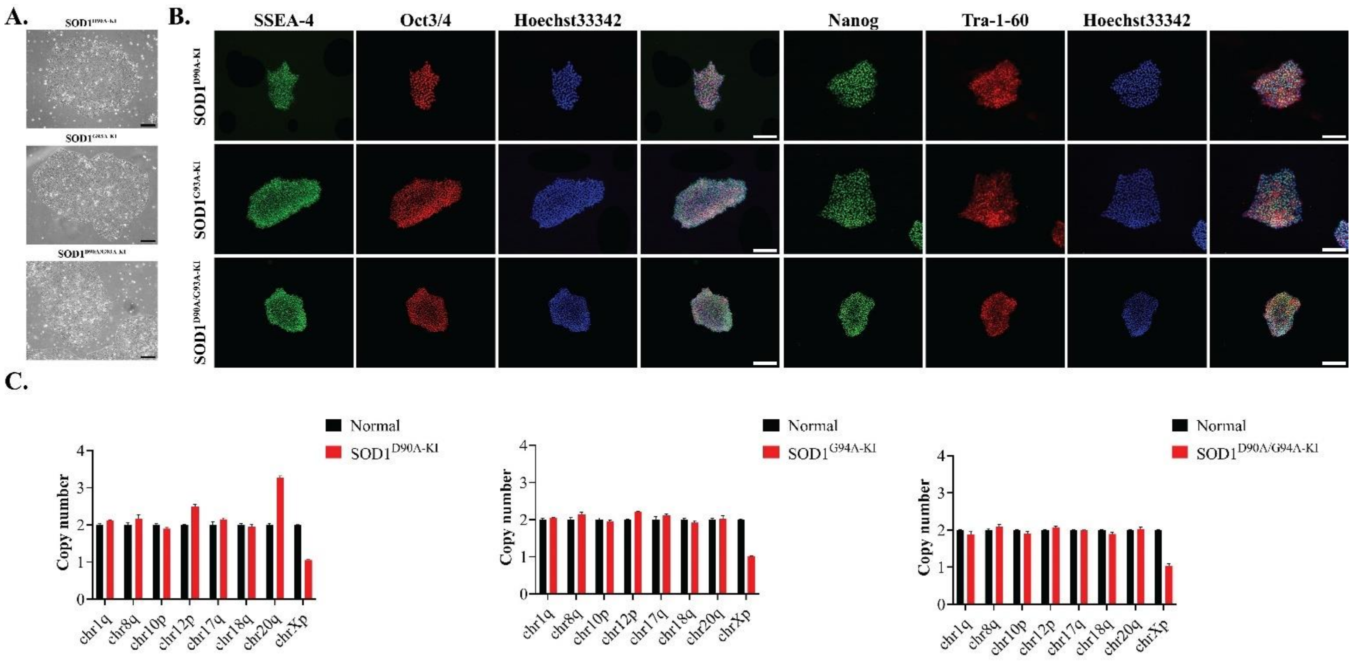


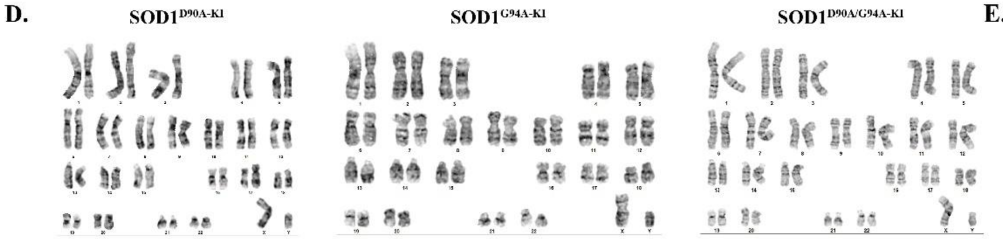


**E.**

| Microbiology/virus screening | |
|---|---|
| Mycoplasma test | Negative |
| Hepatitis B | Negative |
| Hepatitis C | Negative |
| HIV1 or 2 | Negative |

**Supplementary Figure 2. Characterization and quality control of mutant SOD1 iPSC lines. A.** Representative phase-contrast image of $SOD1^{D90A-KI}$, $SOD1^{G93A-KI}$, and $SOD1^{D90A/G93A-KI}$ iPSCs. Scale bar, 100 µm. **B.** Representative images of immunocytochemistry against pluripotency-associated markers Nanog, Tra-1-60, SSEA-4 and Oct3/4. Scale bar, 200 µm. **C.** All iPSC lines have normal chromosome copy number, as assessed by qPCR. Data shown as mean ± SEM, N=3, n=3. **D.** Mutant SOD1 iPSC lines display a normal G-band karyotype. **E.** Mutant SOD1 iPSCs are free from mycoplasma, hepatitis B/C and HIV 1/2 virus.
