## Additional File 10-Supplementary Figure 3 for "Mutations in SOD1 induce ALS-related phenotypes in 3D iPSC-derived motor neuron (MN) spheroids"

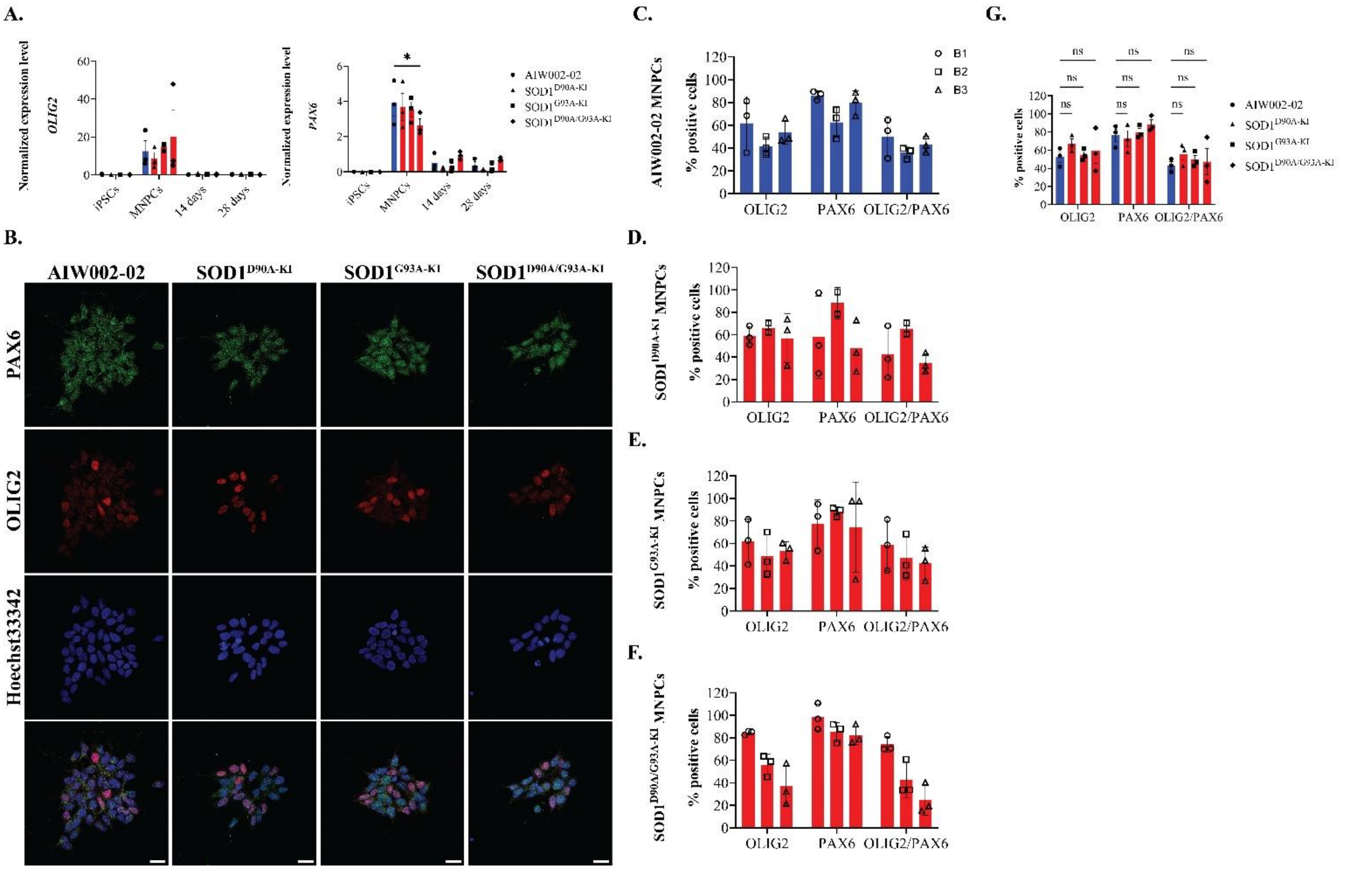

**Supplementary Figure 3. Expression of PAX6 and OLIG2 by MNPCs.** AIW002-02 SOD1$^{D90A\text{-}KI}$, SOD1$^{G93A\text{-}KI}$, and SOD1$^{D90A/G93A\text{-}KI}$ were differentiated into MNPCs as previously described (18, 19, 23, 27). For each cell line, independent differentiation processes were performed to generate three batches of MNPCs, which were stored in liquid nitrogen and later thawed to generate MN spheroids. **A.** At the transcript level, MNPCs expressed OLIG2 and PAX6 as assessed by qPCR. Data shown as mean ± SEM, N=3, n=3. **B-F.** At the protein level, MNPCs were immunostained for Olig2 and Pax6, and the number of Olig2$^{+}$, Pax6$^{+}$, and Olig2$^{+}$/Pax6$^{+}$ cells was quantified. Scale bar, 100µm. Data shown as mean ± SD, n=3. **G.** Statistical analysis performed on the biological triplicates per cell line showed no statistical differences amongst Olig2$^{+}$, Pax6$^{+}$, and Olig2$^{+}$/Pax6$^{+}$ cells. Data shown as mean ± SEM, N=3, n=3. Significance was determined using a one-way ANOVA, followed by a post hoc Dunnett's test using AIW002-02 as the reference sample. *, $p \leq 0.05$; ns, non-significant.
