## Additional File 11-Supplementary Figure 4 for "Mutations in SOD1 induce ALS-related phenotypes in 3D iPSC-derived motor neuron (MN) spheroids"

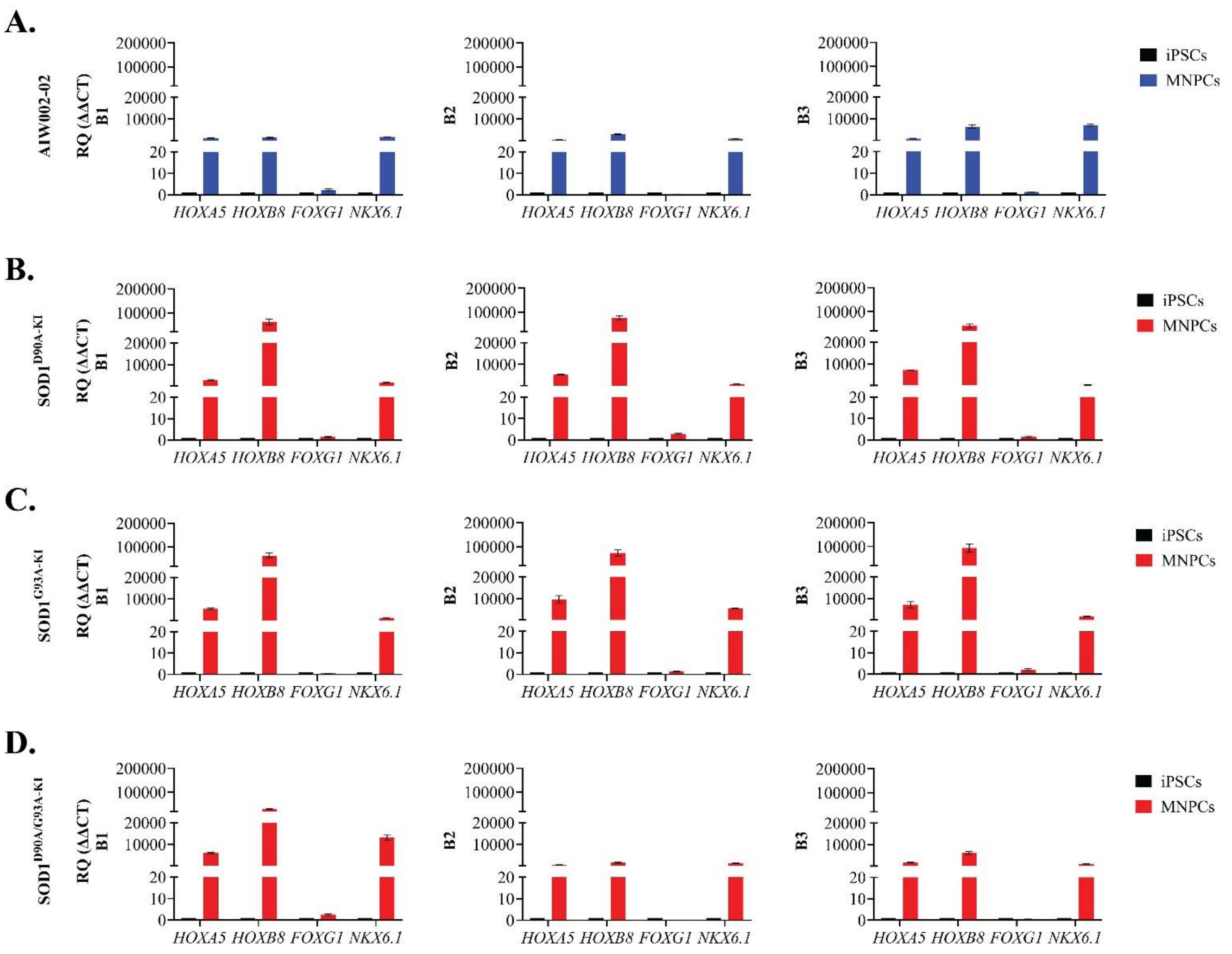


**Supplementary Figure 4. Spatial identity of MNPCs in the spinal cord.** The assessment of HOXA5 and HOXB8 confirmed the spinal lineage of the different MNPC batches induced from the **A.** AIW002-02, **B.** SOD1$^{D90A-KI}$, **C.** SOD1$^{G93A-KI}$, and **D.** SOD1$^{D90A/G93A-KI}$ iPSC lines. FOXG1, a marker of cortical lineage, was used as a negative control. Additionally, the expression of NKX6.1 ascertained the ventral identity of the MNPCs. Bar graphs represent ΔΔCT values, normalized to endogenous controls and compared to iPSCs as the reference sample. Data shown as mean ± SD, n=3.
