## Additional File 12-Supplementary Figure 5 for "Mutations in SOD1 induce ALS-related phenotypes in 3D iPSC-derived motor neuron (MN) spheroids"

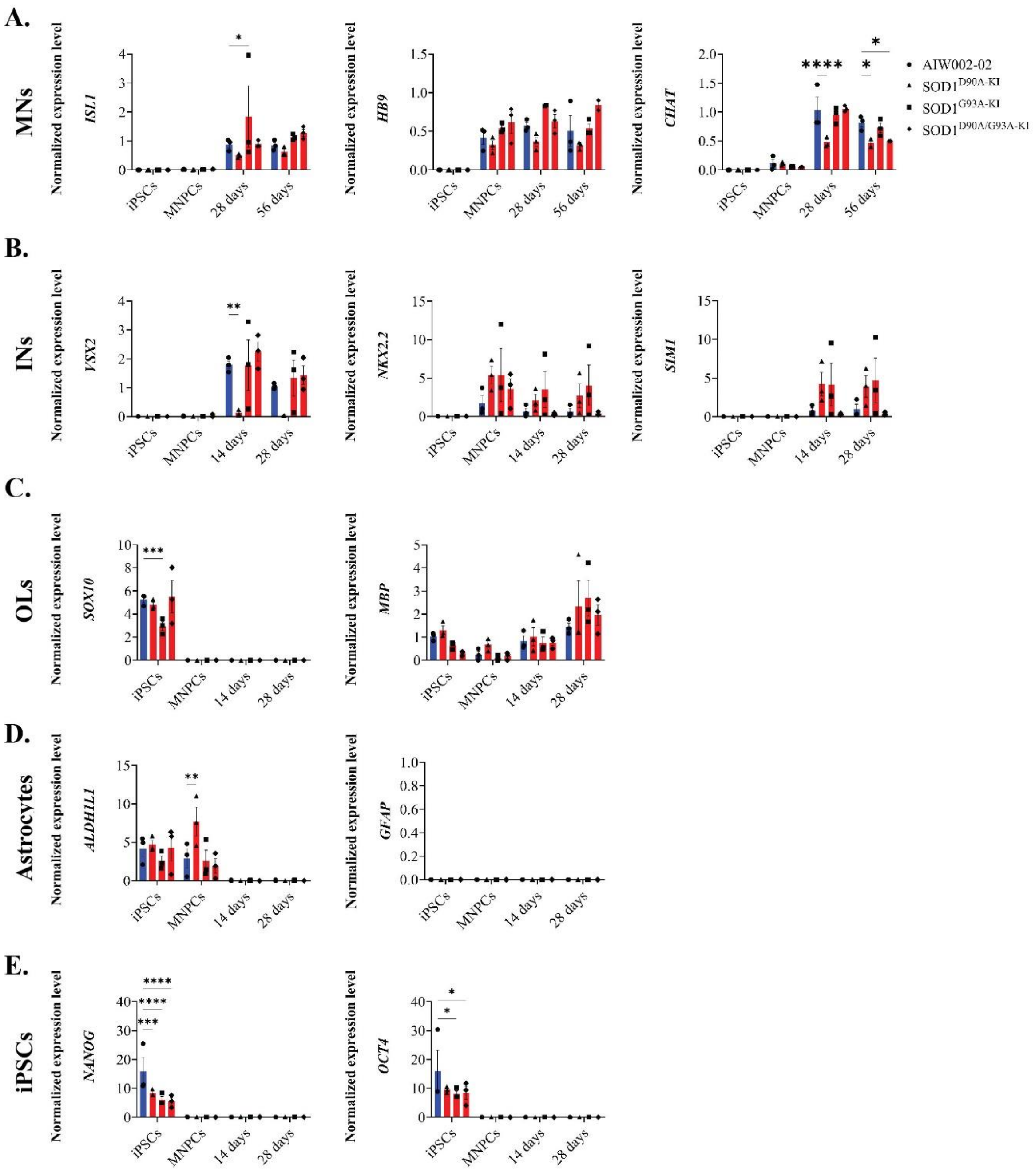


**Supplementary Figure 5. Expression profiling of different cell types within MN spheroids.** The transcriptional expression of markers associated with MNs, interneurons (INs), oligodendrocytes (OLs), astrocytes and iPSCs was analyzed via RT-qPCR across developmental stages, including iPSCs, MNPCs, and MN spheroids at 14, 28, and 56 days. **A.** *ISL1* and *HB9* expression began at the MNPC stage and peaked during the MN spheroid stage, with no significant differences observed between cell lines after 56 days. *CHAT,* a marker of MN maturation, was predominantly expressed in MN spheroids; however, its expression was significantly downregulated in SOD1$^{D90A-KI}$, and SOD1$^{D90A-KI/G93A-KI}$ after 56 days. **B.** Within the dorsoventral axis of the spinal cord, V2 and V3 INs flank the pMN domain. V2 INs were identified by *VSX2* expression, while V3 INs were associated with *NKX2.2* and *SIM1* expression. None of these markers−*VSX2, NKX2.X,* or *SIM1*−showed significant differences across cell lines in MN spheroids after 28 days. **C.** Oligodendrocyte progenitor cells (OPCs) and mature oligodendrocytes were identified through the expression of *SOX10* and *MBP*, respectively. *SOX10* was absent in MN spheroids after 28 days, whereas *MBP* expression was upregulated to comparable levels between cell lines. **D.** **A**strocyte detection was based on *ADLH1L1* and *GFAP* expression. The downregulation of *ALDH1L1* in MN spheroids and the absence of GFAP expression indicate a lack of astrocytes. **E.** As a negative control, the downregulation of the pluripotency-associated markers, *NANOG* and *OCT4,* was evaluated within MN spheroids. Data shown as mean ± SEM, N=3, n=3. Significance was determined using a two-way ANOVA, followed by a post hoc Dunnett's test using AIW002-02 as the reference sample. *, $p \leq 0.05$; **, $p \leq 0.01$; ***, $p \leq 0.001$; ****, $p \leq 0.0001$.
