## Additional File 13-Supplementary Figure 6 for "Mutations in SOD1 induce ALS-related phenotypes in 3D iPSC-derived motor neuron (MN) spheroids"

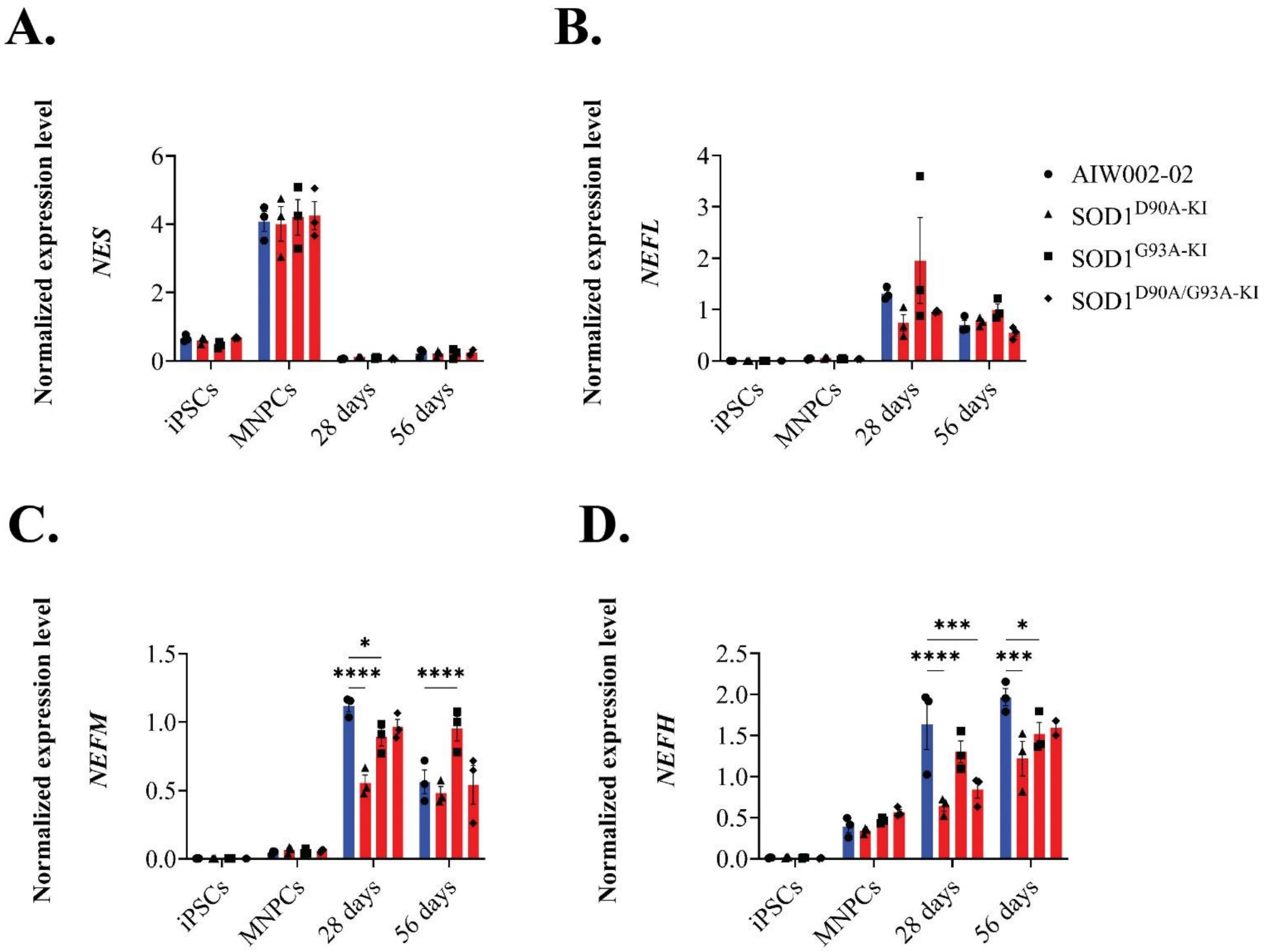


**Supplementary Figure 6. Transcriptional expression of intermediate filaments in MN spheroids.** The expression of **A.** *NES*, **B.** *NEFL*, **C.** *NEFM*, and **D.** *NEFH*, intermediate filaments essential for neuronal function and maturation, was measured through RT-qPCR. Data shown as mean ± SEM, N=3, n=3. Significance was determined using a two-way ANOVA, followed by a post hoc Dunnett's test using AIW002-02 as the reference sample. *, $p \leq 0.05$; ***, $p \leq 0.001$; ****, $p \leq 0.0001$.
