## Additional File 14-Supplementary Figure 7 for "Mutations in SOD1 induce ALS-related phenotypes in 3D iPSC-derived motor neuron (MN) spheroids"

**A.**

| Donor | Sex | Age |
|---|---|---|
| Control 1 | F | 70 y/o |
| Control 2 | M | 55 y/o |
| Sporadic ALS 1 | F | 65 y/o |
| Sporadic ALS 2 | M | 66 y/o |

**B.**

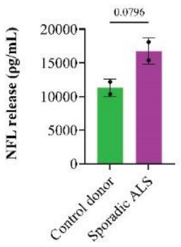


**Supplementary Figure 7. Quantification of NFL release in plasma of control donors and sporadic ALS patients. A.** Plasma samples from two control donors and two sporadic ALS donors were used to validate the ELISA technique used to quantify NFL protein release in the culture media of MN spheroids. **B.** As expected, sporadic ALS donors displayed a trend toward higher levels of NFL protein in plasma than control donors. Data shown as mean ± SD, N=2, n=2. Significance was determined using a t-student.
