## Supplementary figures and images for "Mutations in SOD1 induce ALS-related phenotypes in 3D iPSC-derived motor neuron (MN) spheroids"

### Additional File 15-Supplementary Figure 8

#### Western Blot
##### Gel 1

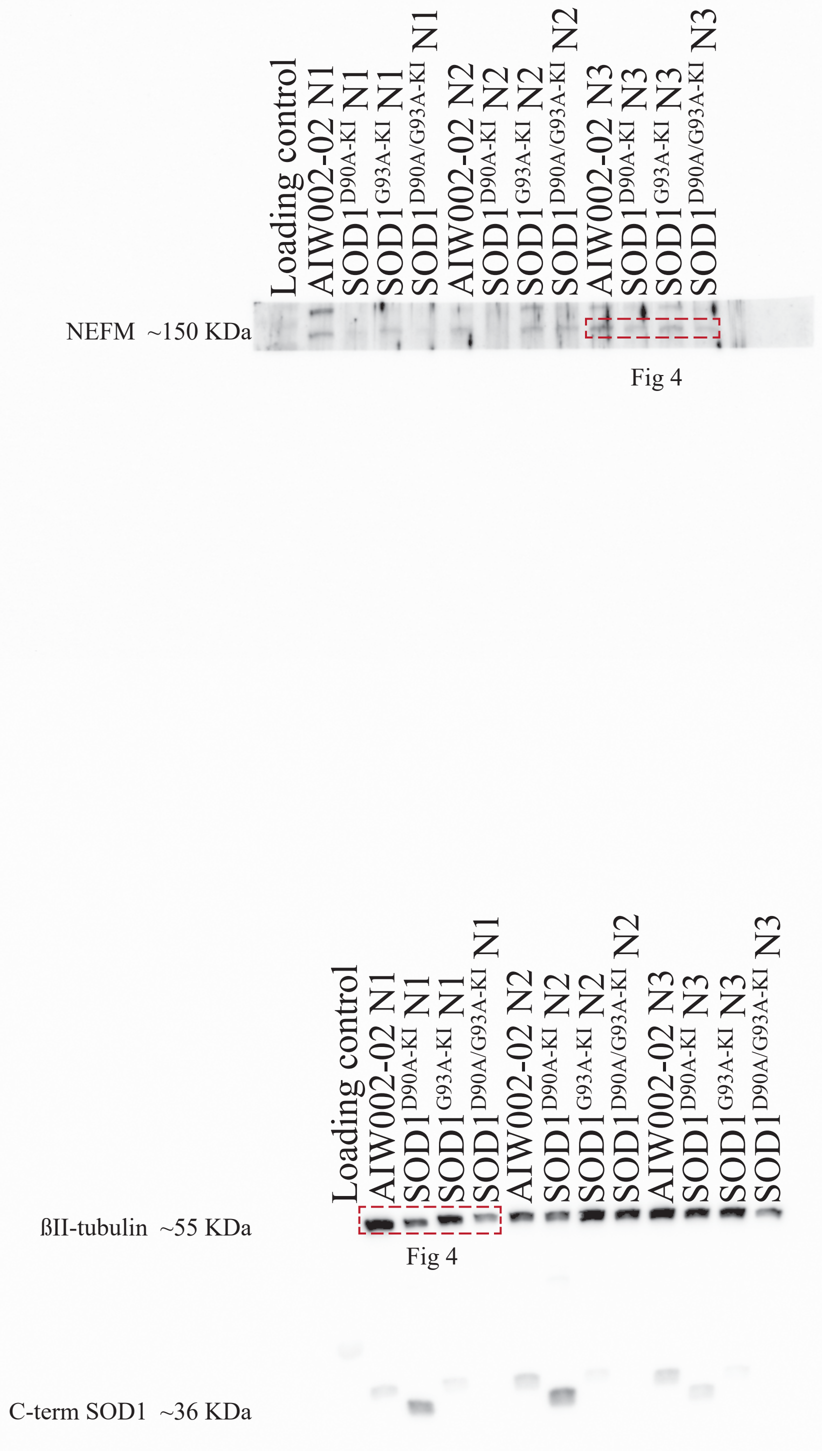

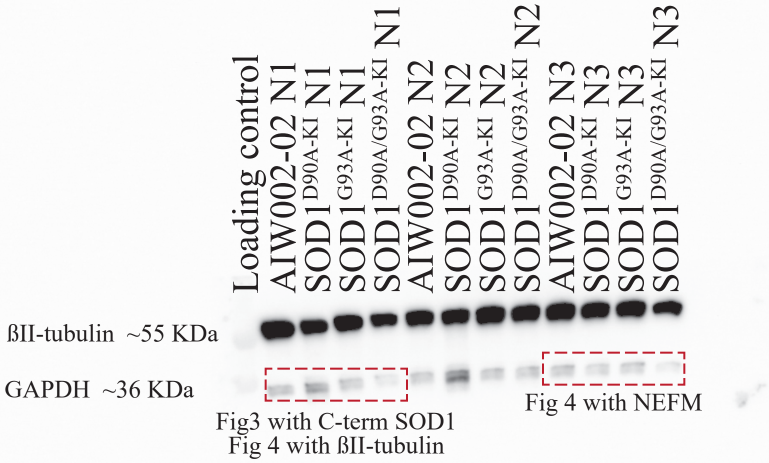

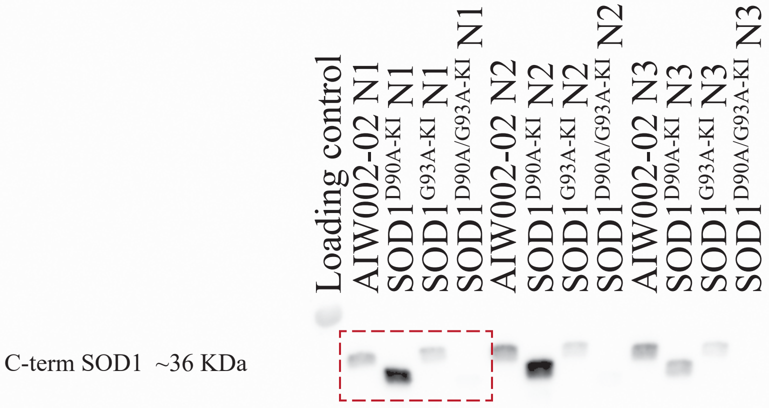

#### Western Blot
##### Gel 2

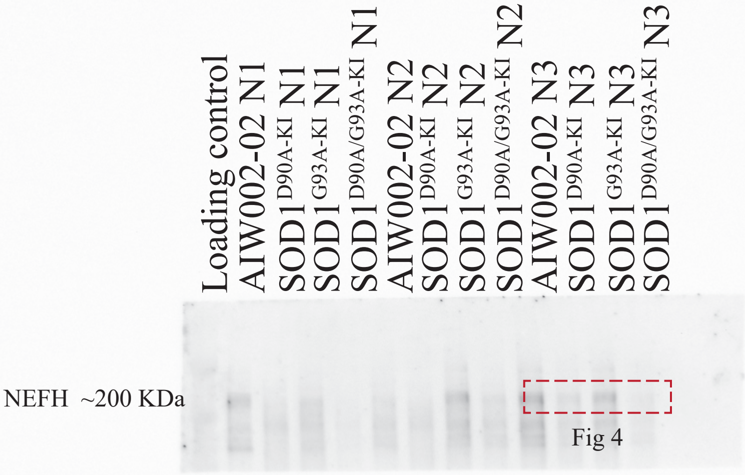


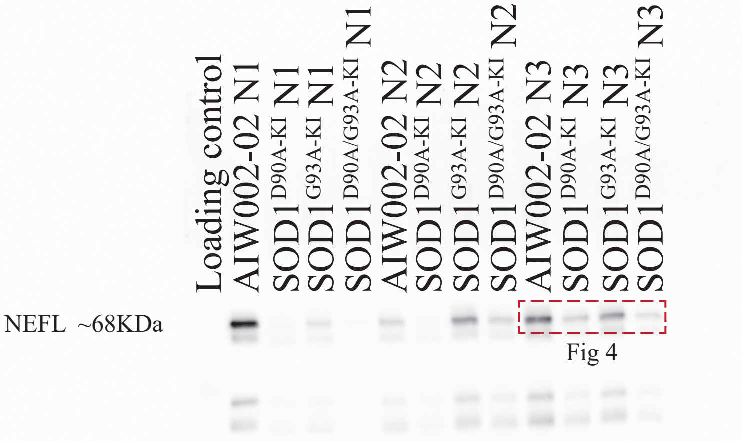

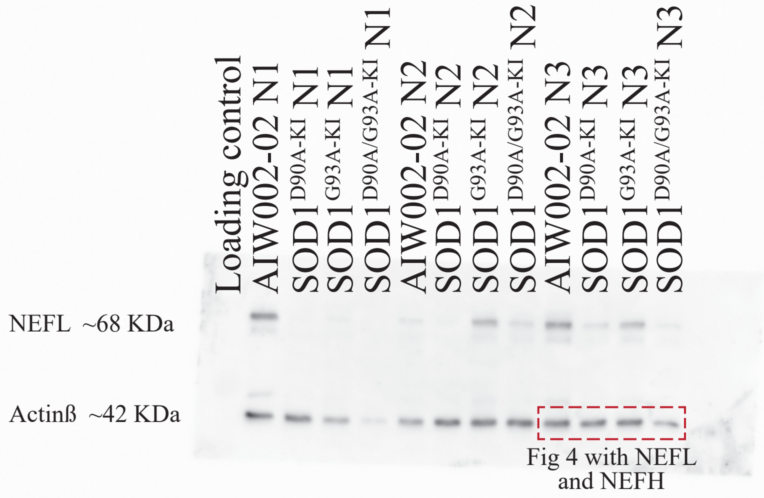

#### Western Blot

##### Gel 3

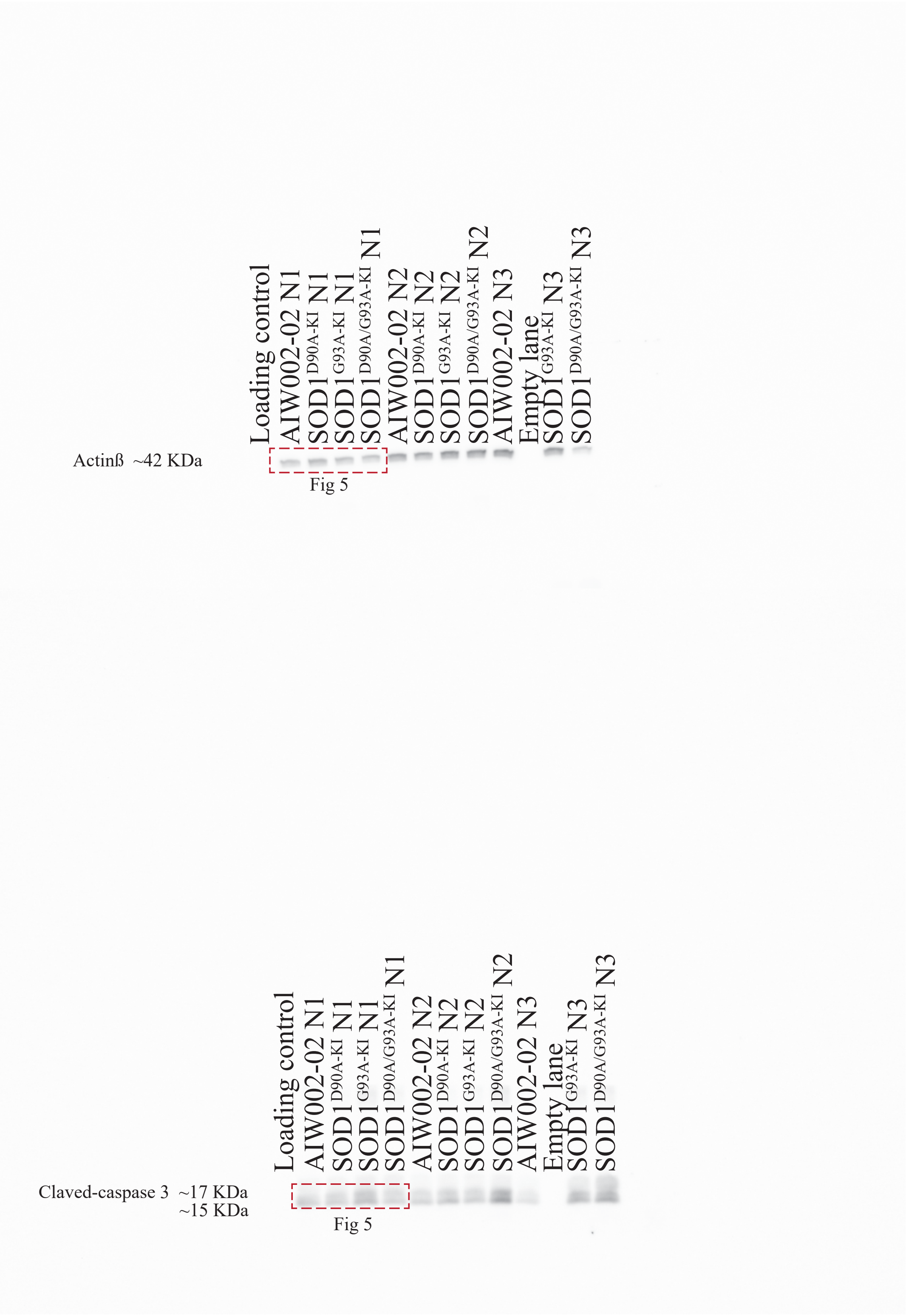
